# BRCA1 and ELK-1 regulate Neural Progenitor Cell Fate in the Optic Tectum in response to Visual Experience in *Xenopus laevis* tadpoles

**DOI:** 10.1101/2021.10.21.465368

**Authors:** Lin-Chien Huang, Haiyan He, Aaron C. Ta, Caroline R. McKeown, Hollis T. Cline

## Abstract

In developing Xenopus tadpoles, the optic tectum begins to receive patterned visual input while visuomotor circuits are still undergoing neurogenesis and circuit assembly. This visual input regulates neural progenitor cell fate decisions such that maintaining tadpoles in the dark increases proliferation, expanding the progenitor pool, while visual stimulation promotes neuronal differentiation. To identify regulators of activity-dependent neural progenitor cell fate, we used RNA-Seq to profile the transcriptomes of proliferating neural progenitor cells and newly-differentiated immature neurons. Out of 1,130 differentially expressed (DE) transcripts, we identified six DE transcription factors which are predicted to regulate the majority of the other DE transcripts. Here we focused on Breast cancer 1 (BRCA1) and the ETS-family transcription factor, ELK-1. BRCA1 is known for its role in cancers, but relatively little is known about its potential role in regulating neural progenitor cell fate. ELK-1 is a multifunctional transcription factor which regulates immediate early gene expression. We investigated the effect of BRCA1 and ELK-1 on activity-regulated neurogenesis in the tadpole visual system using *in vivo* time-lapse imaging to monitor the fate of turbo-GFP-expressing SOX2+ neural progenitor cells in the optic tectum. Our longitudinal *in vivo* imaging analysis shows that knockdown of either BRCA1 or ELK-1 altered the fates of neural progenitor cells, and furthermore that the effects of visual experience on neurogenesis depend on BRCA1 expression, while the effects of visual experience on neuronal differentiation depend on ELK-1 expression. These studies provide insight into the potential mechanisms by which neural activity affects neural progenitor cell fate.

## Introduction

Neurogenesis is the collective process of cell proliferation, differentiation, migration, and survival, which together lead to the generation of mature, functional neurons. During brain development, fate-restricted neural progenitor cells undergo symmetric- or asymmetric-divisions to maintain the progenitor pool, or terminal divisions after which they differentiate into post-mitotic neurons (Gotz and Huttner, 2005). Dysregulation of neurogenesis results in abnormal brain development (Chenn and Walsh, 2002; Gotz et al., 2016; Pramparo et al., 2015). Therefore, insight into the regulation of cell proliferation and neuronal differentiation is essential for advancing our understanding of brain development.

Growing evidence suggests that neuronal circuit activity, driven by spontaneous activity or sensory experience, regulates multiple aspects of brain development (Luhmann et al., 2016; Pan and Monje, 2020). *Xenopus laevis* tadpoles are an excellent experimental system to investigate how sensory experience modulates neural development because tadpoles receive and respond to patterned visual stimuli while neurogenesis and circuit assembly are occurring. In the tadpole, visual stimulation increases the integration of newly generated neurons into the tectal circuit by regulating neuronal structural development, synaptic connectivity and biophysical properties that affect neuronal firing (Aizenman and Cline, 2007; Gambrill et al., 2019; Sharma and Cline, 2010; Sin et al., 2002). Several studies indicate that visual experience also regulates multipotent neural progenitor proliferation, cell fate and neuronal differentiation (Bestman et al., 2012; Sharma and Cline, 2010; Sierra et al., 2015; Song et al., 2012). In a study using CldU to birthdate newly generated cells, exposing animals continuously to dark over 2 days, instead of the normal 12h light/12h dark cycle, significantly increased cell proliferation, expanding the progenitor pool, whereas exposing animals to a simulated motion visual stimulus increased neuronal differentiation (Sharma and Cline, 2010). In an *in vivo* time-lapse imaging study in which neural progenitor cells and their progeny were imaged over time, exposing animals to a simulated motion stimulus for 24h decreased neural progenitor cell proliferation and increased neuronal differentiation, compared to the proliferation and differentiation seen under 12h light/12h dark conditions (Bestman et al., 2012; Sharma and Cline, 2010). These studies indicate that visual stimulation conditions that alter tectal circuit activity lead to different neural progenitor cell fates, however little is known about the molecular mechanisms that regulate these activity-induced cell fate decisions.

Genetic analyses indicate that diverse cellular processes are involved in the regulation of brain development (Hu et al., 2014), suggesting differential analysis of the transcriptomes expressed in neural progenitor cells and newly generated neurons might reveal regulatory pathways involved in neurogenesis. Previous studies have characterized the transcriptomes of neural stem cells from diverse species, and compared the transcriptomes between neural stem cells and neurons (Azim et al., 2015; Berger et al., 2012; van de Leemput et al., 2014), however the potential effects of brain circuit activity on the neural progenitor cell transcriptome *in vivo* have not been reported. In this study, we investigated how different visual experience regimes affect the fate of neural progenitor cells *in vivo* in the optic tectum of tadpoles. We used transcriptomic profiling of neural progenitor cells and newly generated neurons with RNA-seq to identify differentially expressed transcripts. Datamining identified BRCA1 and ELK-1 as candidate molecular regulators of neural progenitor cell fate. We then used *in vivo* time-lapse imaging combined with knock down strategies to demonstrate that the effect of visual experience on the fate of neural progenitor cells depends on ELK-1 and BRCA1 expression and that these regulators play different roles in neurogenesis depending on the visual experience regime.

## Methods

### Animals

Albino *Xenopus laevis* tadpoles of both sexes were obtained from an in-house colony at the Department of Animal Resources at the Scripps Research Institute, La Jolla or purchased from Xenopus Express (Brooksville, FL, USA, RRID:XEP_Xla200). Animals were reared in 0.1X Steinberg’s solution at 22°C under a 12 hour light/12 hour dark cycle, and anesthetized before all procedures in 0.02% tricaine methanesulfonate (MS-222). When experiments were completed, animals were euthanized with 0.1% MS-222. Animals were staged according to Nieuwkoop and Faber (1956). All animal protocols were approved by the Institutional Animal Care and Use Committee of Scripps Research (approval # 08-0083-3).

### Isolation of enriched neural progenitor cells and immature neurons

Cell samples enriched in neural progenitor cells and immature neurons were collected from tadpole midbrain as follows. Stage 46 tadpoles were taken directly from their 12h dark cycle, anaesthetized in 0.02% MS-222, and their brains were electroporated with plasmids expressing turboGFP (tGFP) driven by the oct4/sox2 enhancer from the minimal FGF promoter, called pSOX2-bd::turboGFP, at 2mg/ml, as described (Bestman et al., 2015; Bestman et al., 2012). Immediately following electroporation, animals were exposed to either a simulated motion stimulus, referred to as enhanced visual stimulation, or dark for 24 hours to enrich for tGFP-expressing immature neurons or neural progenitor cells, respectively, as described (Bestman et al., 2015; Bestman et al., 2012; Sharma and Cline, 2010). Midbrains were collected from 100 animals reared in each condition and dissociated into single cells with amphibian PBS (NaCl 113mM, Na2HPO4 8mM, KH2PO4 1.5mM, EDTA 0.1%, EGTA 2mM). Approximately 40,000 tGFP+ cells were collected from ∼100 animals from each condition, using Fluorescence Activated Cell Sorting (FACS; FACSAria II, BD Biosciences, USA; RRID:SCR_018934). For FACS, forward scatter was used to set the threshold for cell size and side scatter was used to set the threshold for cellular granularity. The background fluorescence in the FITC channel was set according to fluorescence from cells dissociated from midbrains without electroporation and cells with green fluorescence higher than the background were collected. Forward and side scatter plots indicated no difference in the size or granularity of the tGFP+ cells compared to the non-electroporated cells, which are a mixture of neural progenitor cells and immature neurons. In addition, there was no difference in size and granularity between tGFP+ neural progenitor cells and immature neurons. In addition, there was no overlap between tGFP+ cells and cells labeled with SytoxRed (S34859, Life Technologies), a nuclear dye that labels unfixed dead cells (data not shown), suggesting that the tGFP+ cells are healthy. Total RNA was extracted using the mirVana kit (Life Technologies, USA), followed by DNase treatment to remove genomic DNA and followed by clean-up using RNeasy mini kit (Qiagen, USA). The samples with RNA integrity number (RIN) >8, measured with a Bioanalyzer were used for subsequent analysis. 2ng of total RNA was amplified to 2–3 µg of double-stranded cDNA, using the Ovation RNA-Seq System V2 (NuGEN, USA). The amplified cDNA was purified, using Agencourt AMPure XP beads (Beckman Coulter, Inc.), quantified by NanoDrop using Agilent Bioanalyzer. Three biological replicates were analyzed for each condition.

### RNA-Seq of neural progenitor cells and immature neurons

1µg of cDNA was sheared in microTube (Covaris) and then used for library preparation (KAPA Taq PCR kits). The size selection for the final PCR product between 200 – 500bp was done by gel purification. The next generation sequencing was done using HiSeq2000 platform (Illumina; RRID:SCR_020132) for single-end reads at size 100bp. Samples were multiplexed in one lane at the Next Generation Sequencing Core at the Scripps Research Institute (La Jolla). Each sample has between 17 and 20 million reads (Supplemental Table S1).

### Bioinformatics analysis

The quality of raw reads was reviewed, using FASTQC (v0.11.4) (http://www.bioinformatics.babraham.ac.uk/projects/fastqc/; RRID:SCR_014583). Two aligners, STAR (v2.4.0j; RRID:SCR_004463) (Dobin et al., 2013) and TopHat2 (v2.0.13; RRID:SCR_013035) (Trapnell et al., 2012), were used to align the reads against the J-strain, v9.1 *Xenopus laevis* genome assembly (Xenbase; RRID:SCR_003280) with the annotation using gene model (JGI v1.8; Xenbase). The v9.1 of the *Xenopus laevis* genome release incorporated >90% of the genome sequence into 18 pseudomolecules representing the 18 chromosomes of *X. laevis*. These 18 pseudomolecules can be categorized as 9 pairs, each pair containing one L and one S pseudomolecule. 45,099 primary transcripts are annotated in v9.1, with primary transcripts being defined as the longest splice variant of a particular gene. Not all the annotated transcripts have a published gene symbol. 17,409 of the transcripts have a published gene symbol; 20,685 have a gene symbol starting with xelaev, which means that this transcript is specific to *X. laevis*; 4,293 start with LOC, which have no published symbols and no orthologs; 2,712 start with xetrop which are homologous to *Xenopus tropicalis* but do not have any published gene symbol. Approximately 38.6% of the primary transcripts have a known gene symbol. These transcripts were used for functional classification with existing databases. For alignment with STAR, no trimming on the raw reads was performed before alignment since STAR itself has a soft clipping function. For TopHat2 alignment, dynamic trimming was performed, using Trimmomatic (v0.32; RRID:SCR_011848) (Bolger et al., 2014). Alignment quality was examined, using SAMStat (v1.09; RRID:SCR_005432) (Lassmann et al., 2011). Only the reads with MAPQ score higher than 20 were included in the differential expression analysis. The reads were counted by HTSeq (HTSeq-count; v0.6.1p1; RRID:SCR_005514; RRID:SCR_011867) (Anders et al., 2015). Differential expression analysis package, DESeq2 (RRID:SCR_015687) (Love et al., 2014), and graphics are performed under R (v3.1.2; cran.r-project.org; RRID:SCR_001905) through Bioconductor (RRID:SCR_006442) (Gentleman et al., 2004). STRING (v10; RRID:SCR_005223) (Szklarczyk et al., 2015) was used for protein-protein interaction analysis; Cytoscape (v3.2.1; RRID:SCR_015784) (http://www.cytoscape.org/) for network analysis and visualization; ClueGO (v2.1.7; RRID:SCR_005748) (Bindea et al., 2009) for functional network analysis; PANTHER (RRID: SCR_004869) (Mi et al., 2013) for Gene Ontology (GO) analysis; ENCODE; RRID:SCR_015482) (Consortium, 2012) for transcription factor analysis. Fragment Per Kilobase of transcript per Million mapped reads (FPKM) was calculated, using Cufflinks suite (v2.2.1; RRID:SCR_014597) (Trapnell et al., 2012).

### *In vivo* time-lapse imaging

For *in vivo* live-cell time-lapse imaging, whole-brain electroporation was performed on late stage 46 tadpoles with 2 μg/μl pSOX2-bd::turboGFP (Bestman et al., 2015) and 0.4mM antisense morpholino oligonucleotide tagged with lissamine fluorophores, against *brca1* transcript (GGTTCCATTTGTGTCAGCTCTCAGC) and against *elk-1* transcript (GGTCATTTTACTTTGTCCTGTCCCT), or a control non-specific sequence (GeneTools, Philomath, OR). Animals were divided into two groups for the 3-day duration of the time-lapse imaging; one group housed in the control 12h light/12h dark condition and the other housed in the dark in a light impermeable black chamber. Animals were maintained in their respective housing conditions throughout the experiment except when imaging. Tadpoles were screened for consistent morpholino fluorescent labeling in the tectum. For imaging, tadpoles were anesthetized in 0.01% MS-222, placed in a custom-built chamber and imaged with 20X (Olympus XLUMPlanFL 0.95 NA) water immersion lens on a custom-built two-photon microscope modified from an Olympus FV300 system (Bestman and Cline, 2008; Ruthazer et al., 2006). A stack of images for each tectal lobe was acquired at 1µm intervals ranging from 120µm to 160µm depending on the distribution of tGFP+ tectal cells over three days. All samples were imaged in parallel using identical image acquisition parameters. Analysis was conducted using Cell Counter plugin in FIJI, an image processing package of ImageJ (RRID:SCR_002285) (Schindelin et al., 2012; Schneider et al., 2012). tGFP-labelled cells were identified and categorized into three groups, neural progenitor cells, neurons or unidentifiable based on their morphology, using criteria as described (Bestman et al., 2015; Bestman et al., 2012). While the majority of tGFP+ cells could be classified as either neural progenitor cells or neurons, a small population of cells was unclassified or unidentifiable. In control groups, the fraction of unidentifiable cells was 0.5-4.4% across all experiments over the course of the 3 days. For BRCA1 KD, the unidentifiable cell population ranged from 0.4-5.9%, and for ELK-1 the fraction of unidentifiable cells was 3.1-10.8% across all experiments and timepoints. The proliferation and survival rates were calculated based on the changes in total number of tGFP-labelled cells at day 2 and day 3 normalized against the number at day 1. The changes in the fate of neural progenitor cells as the result of the treatment is presented as the percentage of each cell type comprised the total cell population each day. For experiments testing for interactions between the effects of visual experience and BRCA1 or ELK-1 expression on neural progenitor cell fate, we used a factorial experimental design (Collins, 2018) which allows us to compare results between multiple experimental conditions.

### Immunohistochemistry

Animals were anesthetized and then fixed with freshly made 4% paraformaldehyde (Electron Microscopy Sciences, Fort Washington, PA) in 1x phosphate-buffered saline (PBS; pH 7.4) with a brief microwave pulse (150mV on-off-on, 1min each; Pelco BioWave Pro microwave, Model 36500, Ted Pella, Redding, CA) and were post-fixed at 4°C overnight. Whole brains were dissected and incubated in blocking solution (5% normal donkey serum and 1% Bovine Serum Albumin (BSA; Sigma) in PBS with 0.1% Triton-X100 (PBS-T)) for 1 hour at room temperature before transferred to the anti-pH3 antibody solution (1:200 in blocking solution; #9706, Cell Signaling; RRID:AB_331748) at 4°C for 3 days. After washes with PBS-T, brain tissues were incubated in secondary antibody solution (Alexa488 donkey anti-mouse secondary antibody, 1:1,000; A21202, Life Technologies; RRID:AB_141607) at 4°C overnight. After PBS-T washes, for cell death analysis in fixed tissue, nuclear labeling using Sytox Orange (SytoxO, 1:5000; S11368, Life Technologies) was applied to the brain tissues for 20 mins at room temperature (Faulkner et al., 2015). After several washes with PBS-T, brain tissues were mounted in 6M urea in 50% glycerol for imaging. 36µm Z-series we collected at 1µm intervals using Nikon C2 (20x Plan Apo lens with 0.75 NA), and ImageJ Cell Count plugin was used for analysis. Apoptotic cells were identified based on the morphology of small granular structure and the high labeling intensity (Thompson and Cline, 2016), and categorized into two groups, neural progenitor cells (NPC) and neurons based on their location in the tectum.

### Western blots

To test the effect of different visual experience conditions, animals were reared under enhanced visual stimulation or dark for 30 hours as described previously (Sharma and Cline, 2010) and the midbrains were dissected for homogenization. To test the effect of morpholino knockdown, whole-brain electroporation was performed at late stage 46 tadpoles with 0.4mM morpholinos, and the midbrain tissues were dissected 2 days later. Tissues were homogenized in different lysis buffers for different antibodies. Experimental and paired control samples were prepared and processed side by side. For ELK-1 (Abcam #ab188316; RRID:AB_2890919) and SOX2 (Cell Signaling Technology #3579S; RRID:AB_2195767) antibodies, tissues were homogenized in RIPA buffer with brief sonication, and concentration was measured by BCA assay. 10µg of lysate was run on a Mini-Protean TGX precast gels (BioRad). For BRCA1 antibody (SCBT #SC-646; RRID:AB_630945), tissue was homogenized in lysis buffer documented in (Joukov et al., 2001); HEPES 100mM (pH 7.5), NaCl 200mM, EDTA 40mM, EGTA 4mM, NaF 100mM, β-glycerophosphate 20mM, sodium orthovanadate 2mM, Nonidet P-40 1%, Complete Protease Inhibitor mixture 1:50), and concentration measured by DC Protein Assay (BioRad). 40mg of lysate was loaded onto in-house made 7% gel. Proteins were transferred to a nitrocellulose membrane and blotted with standard protocols. Antibodies were detected by goat anti-mouse/rabbit HRP-conjugated secondaries (BioRad) followed by ECL (Pierce,Thermo Fisher Scientific, 32209). Quantification was performed using densitometry (ImageJ), different exposures were used to avoid saturation, and bands were normalized to total protein using Ponceau S (Romero-Calvo et al., 2010).

### Statistical tests

The nonparametric Mann-Whitney and one-tailed or two-tailed Student’s t test were used for comparisons of two groups using Prism 9 statistics software (Graphpad Prism; RRID:SCR_002798). For unbalanced two-way ANOVA analysis, car package in R was used (R Core Team, 2018; RRID:SCR_001905).

### Data Availability

All raw *X. laevis* data are available on GEO as [pending] and on Xenbase. pSox2-bd::FP plasmid is available from Addgene, plasmid #34703. Morpholino sequences are provided in the accompanying reagent list. The differential expression read data are provided in the supplemental material as Supplementary Data S1. The Panther analysis, STRING analysis, and CytoScape data are provided in the supplemental material as Supplementary Data S2, S3, and S4, respectively.

## Results

### Isolation of neural progenitor cells and immature neurons from the optic tectum

Our previous studies indicate that maintaining Xenopus tadpoles in the dark increases neural progenitor cell proliferation in the optic tectum, expanding the progenitor pool, and in contrast exposing tadpoles to visual stimulation increases the differentiation of progenitors into neurons (Bestman et al., 2012; Sharma and Cline, 2010), suggesting that activity-dependent molecular changes in neural progenitor cells might affect neural progenitor cell fate. To identify transcripts that mediate these visual experience-induced changes in neural progenitor cell fate, we conducted a transcriptome analysis using RNA-seq to profile transcripts in neural progenitor cells and their neuronal progeny in animals exposed to different visual experience regimes. To isolate enriched populations of neural progenitor cells and immature neurons, we expressed tGFP in neural progenitor cells *in vivo* by electroporating the optic tectum with a plasmid that drives tGFP expression upon binding of endogenous SOX2 (Bestman et al., 2015; Bestman et al., 2012). *In vivo* imaging indicates that tGFP+ neural progenitor cells and their newly differentiated neuronal progeny can be identified within 24 hours after electroporation (Bestman et al., 2012). We then exposed animals either to dark or visual experience using a simulated motion stimulus to bias the fate of tGFP+ neural progenitors toward proliferation, generating more neural progenitor cells or toward neuronal differentiation, respectively (Figure 1A, B). Next, we dissected midbrains from animals exposed to dark or visual experience and isolated approximately 40,000 tGFP+ neural progenitor cells and immature neurons by fluorescence-activated cell sorting (FACS), based on their tGFP fluorescence compared to non-electroporated cells (Figure 1C). We observed no overlap between tGFP+ cells and cells labeled with SytoxRed, a nuclear dye that labels dead cells, indicating that the tGFP+ cells are healthy (data not shown). The quality of total RNA isolated from these sorted cells has high integrity (Supplemental Figure S1, A), with RIN values of 9.4 for neural progenitor cells and 9.2 for immature neurons (data not shown). Taken together, these data validate our protocol for isolating enriched populations of neural progenitor cells and immature neurons.

**Figure 1.**
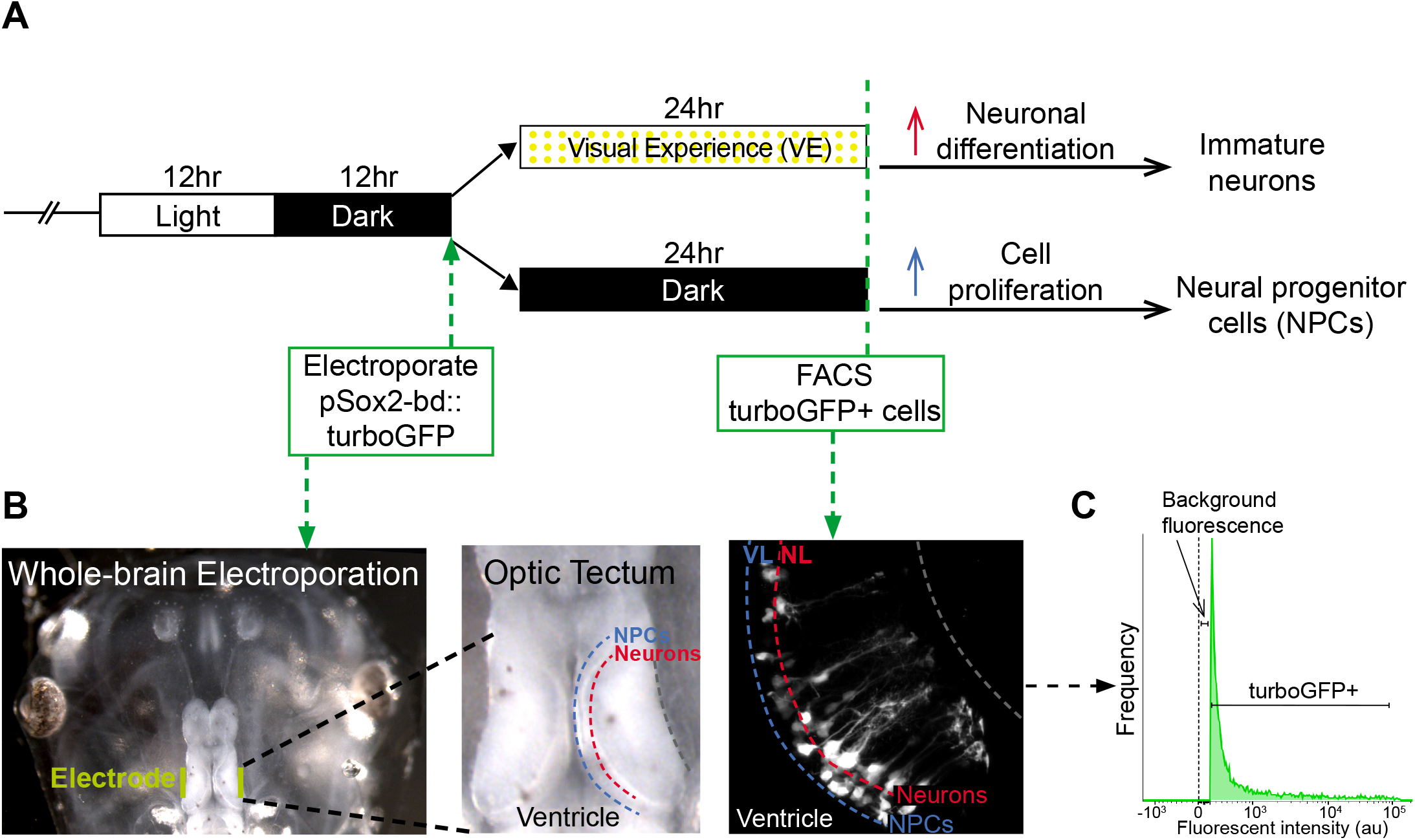
Protocol to generate neural progenitor cells and immature neurons in optic tectum of *Xenopus laevis* to isolate RNA. **A.** Visual experience paradigm used to enrich for neural progenitor cells and immature neurons. Animals are reared in 12h light/12h dark until stage 46 when the midbrain is electroporated with pSOX2-bd::turboGFP plasmid. After electroporation, animals are reared in conditions of either enhanced visual experience (VE) for 24 hours to induce neuronal differentiation and enrich for immature neurons, or visual deprivation (dark) to drive cell proliferation and enrich for neural progenitor cells (NPCs). **B.** *In vivo* tGFP labeling cells in the tadpole midbrain by whole brain electroporation. pSOX2-bd::turboGFP (tGFP) in injected into the ventricle and electroporated into the brain with electrodes positioned next to the mid-brain region (red). After 24h, tGFP is detected in neural progenitor cells in the ventricular layer (VL, blue) and immature neurons in the neuronal layers (NL, red). From each visual condition, midbrains were isolated, cells dissociated, and tGFP+ cells sorted by FACS. **C.** Fluorescence histogram demonstrates the gate setting of the fluorescence-activated cell sorting (FACS) to isolate tGFP+ cells. Control is set to the background fluorescence from non-electroporated midbrain cells.

### Accessing RNA-seq read alignment against *Xenopus laevis* genome scaffold v9.1

To identify transcripts that might affect cell proliferation or neuronal differentiation, we characterized the differences in the transcriptomes of the neural progenitor cell and immature neuron cell populations. We profiled the transcript expression using v9.1 of the *Xenopus laevis* genome assembly and annotation (JGI gene model v1.8) for alignment. Approximately 39% of the transcripts have a known gene symbol, and only these transcripts were used for functional classification with the existing databases.

In each of our 3 biological replicates, there are approximately 17-20 million single-end reads at 100bp (Supplemental Table S1), and the reads were determined to be high-quality by FASTQC (Supplemental Figure S1A). To include all the potential changes in transcript expression, two aligners or mappers were used, Spliced Transcripts Alignment to a Reference (STAR) and TopHat2. Each aligner, due to the nature of different algorithms they incorporate, has different true positive rates and false positive rates. The high quality of alignment was summarized by SAMStat with 92.9% of the reads having a MAPQ score ≥30 (Supplemental Figure S1 B). Including the output from both aligners provides a more inclusive picture of the transcriptome analysis. The efficiency of alignment against the genome assembly is 80% and 78% on average among all the samples, using STAR and TopHat2, respectively (Supplemental Figure S1, C-H). The reads with an alignment score < 20 were not included in the differential expression analysis, to ensure the specificity of the read count for each transcript. Over 14 million reads on average were uniquely aligned to the genome, which provided enough depth in the sequencing for a reliable differential expression analysis. We further characterized where these reads were aligned using STAR and found that 64% of the reads on average were aligned to transcript regions annotated on the genome scaffolds, while 35% of the reads on average were aligned to intergenic regions (Supplemental Figure S1, C-E). The alignment of 1% of the reads could not be determined since they were aligned to multiple transcripts. In a similar analysis using TopHat2 as the aligner, 62% of the aligned reads were assigned to a transcript; 30% aligned to intergenic regions and 8% were classified as ambiguous (Supplemental Figure S1, F-H). There was no apparent difference in the percent of aligned reads against intergenic region between neural progenitor cells and immature neurons. To examine where the reads are aligned in the transcripts, we categorized the transcript alignment into different regions: 56% of the reads align to coding domains (CDS), 16% to introns, 4% to 5’UTR, and 24% to 3’UTR using STAR (Supplemental Figure S1 E), and 55% CDS, 16% introns, 8% 5’UTR, and 21% 3’UTR using TopHat2 (Supplemental Figure S1H). Taken together, these data indicate the high quality of our reads and read alignments to the *Xenopus* genome and demonstrate the distribution of read alignments in the coding and non-coding regions. High quality of reads and read alignments are essential for a reliable differential expression analysis and subsequent analysis.

### Differential expression analysis of transcripts expressed by neural progenitor cells and immature neurons

We next profiled the differences in neural progenitor cell and immature neuron transcriptomes and identified the transcripts that are enriched in one cell population or the other, by conducting differential expression analysis on transcript expression with DESeq2 on the alignment output from STAR and TopHat2. Out of 45,099 transcripts, 27,027 and 26,137 transcripts were detected, using aligner STAR and TopHat2, respectively, based on the criteria that the total number of normalized counts is larger than 6 across 6 samples, i.e. one count per sample on average. The difference in the number of transcripts detected between STAR and TopHat2 likely results from a difference in the number of reads that are classified as ambiguous. We used two ways to detect changes in transcript expression: one calculates differential expression based on the reads aligned to the whole transcript/mRNA, while the other way calculates the reads aligned to the CDS only. The statistical analysis based on CDS only shows which transcripts are differentially expressed between neural progenitor cells and immature neurons and have the potential to be translated into proteins, while the analysis based on mRNAs incorporates the information from 5’UTR, 3’UTR and introns in addition to CDS. We identified 487 or 464 differentially expressed transcripts based on the mRNAs and 738 or 677 differentially expressed transcripts based on the CDS using STAR or TopHat2, respectively, as the aligner. The statistical significance was determined using a false discovery rate less than 0.1 and based on a requirement for a minimal fold change of 4 (log2(2.0)) for the transcript expression to be called as differentially expressed between cell populations. To examine how closely correlated the output is from two different aligners, we compared the fold change calculated based on the output from STAR vs TopHat2. The fold change of the transcript expression between STAR and TopHat2 is well correlated in a linear relationship. In addition, 67.4% (383/568) of differentially expressed transcripts, calculated based on the mRNAs (Supplemental Figure S2 A) and 71.7% (591/824) of differentially expressed transcripts based on CDS (Supplemental Figure S2 B) were overlapped between two aligners. The high percentage of transcripts found differentially expressed in the analyses based on both aligners further supports a strong correlation between the respective outputs from STAR and TopHat2. To further assess whether the analyses performed based on mRNA and CDS, using the same aligner, are well-correlated, a comparison between fold change of transcript expression was performed. A linear relationship was observed in the fold change of the transcript expression between CDS and mRNA, using STAR (Supplemental Figure S2 C). In addition, an MA plot of the mean expression and fold change of the transcript expression between neural progenitor cells and immature neurons based on the CDS, using STAR, shows that the mean expression (the number of normalized reads) of differentially expressed transcripts ranged from low to medium expression level (Supplemental Figure S2 D), indicating the analysis is unbiased based on expression level. From these data, we identified a total of 1,130 transcripts that were differentially expressed between neural progenitor cells and immature neurons from both aligners based on mRNA and CDS (Supplemental Data S1).

To validate the specificity of the enrichment of neural progenitor cells and immature neurons from our dataset, we tested for the differential expression of genes that are known to be involved in neuronal differentiation or proliferation. We found that the immature neuron population has higher expression of transcripts involved in neural patterning and differentiation, such as *neurod1, wnt1, fgf2, vegfa, nfkb1,* and *smad9* (Table 1). Conversely, neural progenitor cells expressed transcripts known to be involved in proliferation, including e*lk-1, e4f1, sstr4, bmp4, jak2* and *nr2f5* (Table 1). Taken together, the differential expression analysis of the transcripts recovered from cell populations enriched in neural progenitor cells and immature neurons demonstrated a well-correlated fold change in transcript expression between 2 aligners, TopHat2 and STAR. In addition, the enrichment procedure for the neural progenitor cells and immature neuron populations was confirmed by expression of known neuronal and progenitor cell markers in the RNA-seq data. Based on these analyses, we then used the list of 1,130 differentially expressed transcripts, which includes transcripts identified by either STAR or TopHat2 aligners, for further functional classification.

**Table 1.** Canonical progenitor and neuronal genes are enriched in isolated neural progenitor and neuronal samples.

| Enriched in<br>Neural Progenitor Cells | Fold Change<br>(log<sub>2</sub>) | Enriched in<br>Immature Neurons | Fold Change<br>(log<sub>2</sub>) |
| --- | --- | --- | --- |
| elk1 | 6.4 | neurod1 | 8.0 |
| e4f1 | 6.3 | wnt1 | 7.6 |
| sstr4 | 5.9 | fgf2 | 6.8 |
| bmp4 | 4.8 | vegfa | 6.2 |
| jak2 | 3.5 | nfkb1 | 4.5 |
| nr2f5 | 3.0 | smad9 | 3.5 |

### PANTHER protein class analysis identified functional categories of differentially expressed genes

To investigate the involvement of the differentially expressed transcripts in cell proliferation and neuronal differentiation, we functionally categorized these transcripts using the PANTHER (Protein Analysis Through Evolutionary Relationships) classification system. Of the 1,130 transcripts that were differentially expressed between neural progenitor cells and immature neurons, 635 were annotated with a published gene symbol in the genome assembly (v9.1), which then could be used for functional analysis. 630 out of the 635 differentially expressed transcripts were recognized by the PANTHER database, and 367 out of 630 genes were clustered based on PANTHER protein classification. The PANTHER protein classification categorized more genes than any other classification scheme in the PANTHER system, such as GO slim biological process, cellular component and molecular function, and PATHWAY analysis. The 367 PANTHER-classified transcripts were clustered into 4 major protein categories: catalytic activity (220 genes), DNA binding (98), receptor-mediated signaling (146) and structural proteins (89) (Figure 2A, Supplemental Data S2). The GO protein classes with the most differentially-expressed genes were nucleic acid binding (52), enzyme modulator (47), and transcription factor (46). These results indicate that catalytic activities and transcription are heavily involved in either maintaining self-renewal capacity or neuronal differentiation.

**Figure 2.**
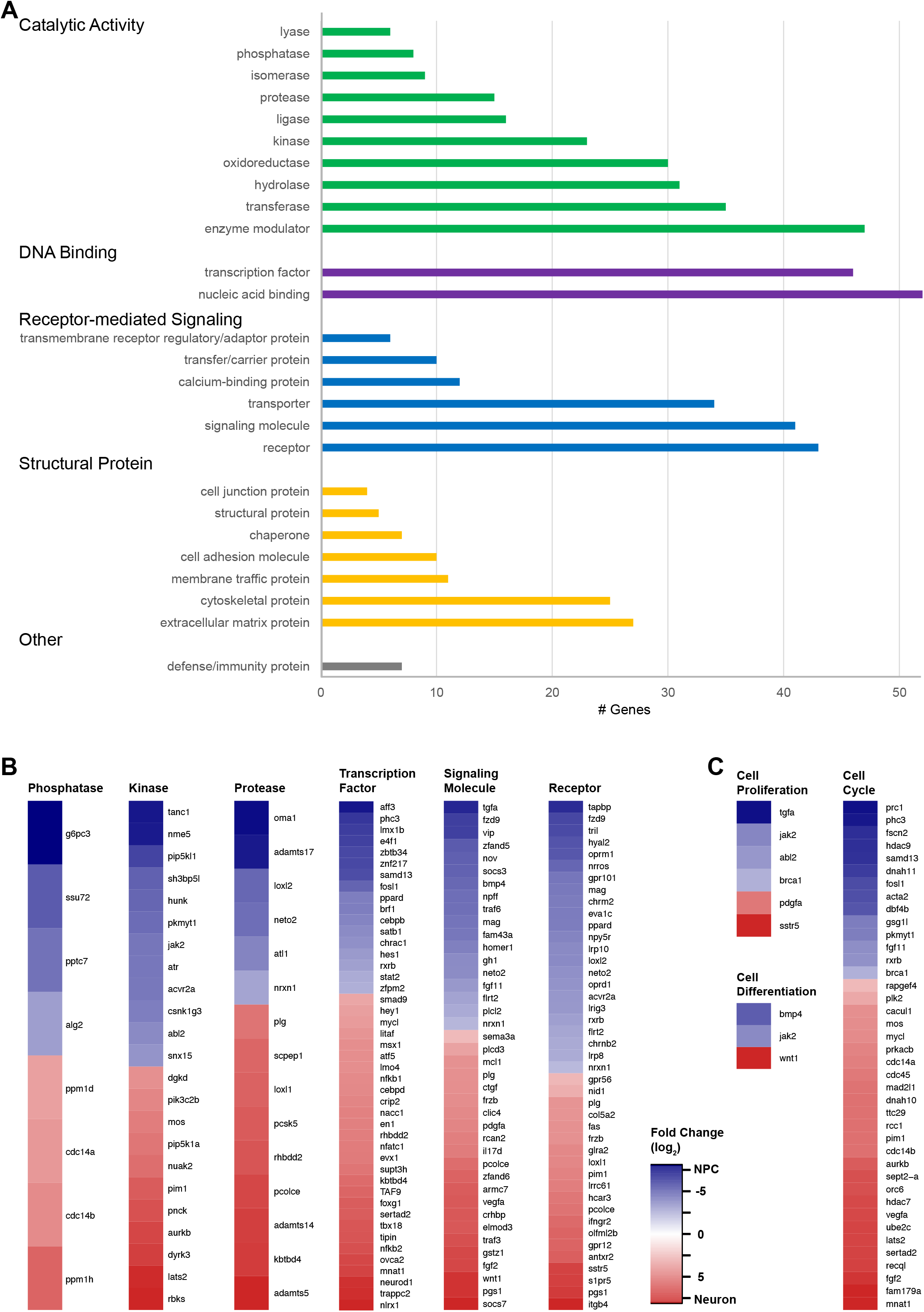
Gene Ontology (GO) analysis of differentially expressed transcripts. **A.** Differentially expressed transcripts are categorized based on GO protein classes. Gene number per GO class is indicated on the X-axis. **B.** Fold change of transcript expression between neural progenitor cells and immature neurons in selected protein classes. **C.** Fold change of transcript expression between neural progenitor cells and immature neurons in GO biological processes: cell proliferation, cell differentiation, and cell cycle.

Kinases and phosphatases, proteases, receptors and signaling molecules, and transcription factors were enriched in our differential expression dataset (Figure 2B). Further analysis of these subcategories indicated that there was no obvious bias in the direction of transcript expression fold changes between neural progenitor cells and immature neurons. PANTHER identified 6 proteases that were enriched in neural progenitor cells, and 9 that were enriched in immature neurons (Figure 2B). In addition, 45 transcription factors were differentially expressed between neural progenitor cells and neurons (Figure 2B, Supplemental Data S2. Transcription factors enriched in neural progenitor cells include *e4f1, znf217* and *limx1b*, while those enriched in neurons include *neuroD1, foxg1* and *evx1* (Figure 2B). Three GO slim biological processes, cell proliferation, cell differentiation and cell cycle were also identified by PANTHER as enriched in our differential expression data (Figure 2C, Supplemental Data S2). In the cell cycle category, *brca1, tgfa* and *jak2* were enriched in neural progenitor cells, while *sstr5* and *pdgfa* expression were relatively enriched in neurons (Figure 2C, right). Together, these analyses demonstrate that PANTHER’s categorization of 367 differentially expressed transcripts between neural progenitor cells and immature neurons provides a broad picture of the differential expression patterns of these genes.

### Protein interaction networks identify hub proteins: key players in neurodevelopment among differentially expressed genes

Protein function is often inferred by protein-protein interactions. To better understand the potential functions of the protein products of the differentially expressed genes, we performed a network analysis of protein-protein interactions using STRING and Cytoscape (Figure 3A,B). STRING is a database of known and predicted protein interactions based on bioinformatic analysis of genomic and proteomic data and on published work. Starting again from the 1,130 differentially expressed transcripts, of which 635 have a published gene symbol, 629 of the 635 were recognized by the STRING protein database. Of the 629 proteins predicted from the annotated transcripts, STRING identified 458 proteins as having one or more interaction partners (Supplemental Data S3). We graphed the number of protein-protein interactions versus the degree of differential expression of the transcripts, considering degree centrality and closeness centrality as two key metrics of the importance of nodes within networks (Figure 3A)(Wang and Zhao, 2015). Degree centrality in the network reflects the number of interactions (i.e., degree) each protein has within a node. Closeness centrality is a measure of the shortest path between proteins within a node. In this case, a high closeness score indicates that the interactions of a protein and its neighboring partner protein are less likely to be by-passed by other proteins in the node. This means that the protein plays an irreplaceable role in the network. Ranking protein importance based on degree centrality and closeness centrality (Wang and Zhao, 2015) indicates that proteins with degree centrality of greater than 20 are likely to play an important role in neurogenesis (Figure 3A). Nine proteins had 20 or more interaction partners, and 4 of these, ACTA2, BMP4, JAK2 and BRCA1, were enriched in neural progenitor cells (Figure 3A, blue), while the other 5, ITGA2, VEGFA, FGF2, AURKB and NFKB1, were enriched in immature neurons (Figure 3A, red). Network analysis indicated that these 9 proteins interact with each other (Figure 3B).

**Figure 3.**
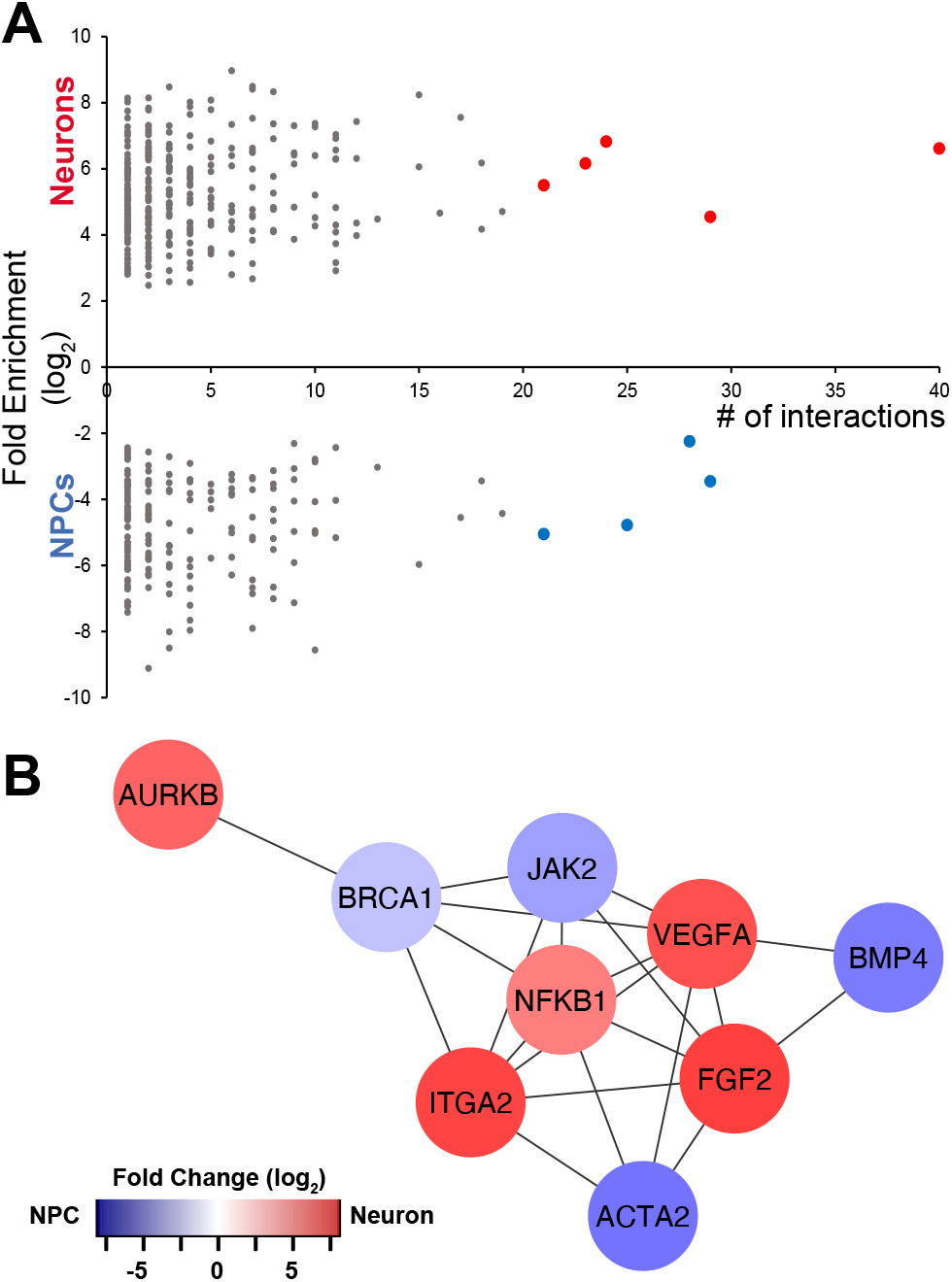
Functional network analysis for the differentially expressed transcripts between neural progenitor cells and immature neurons. **A.** Protein interaction network analysis of differentially expressed transcripts, arranged by number of binding partners (degree). Blue transcripts indicate the most enriched in neural progenitor cells; red transcripts are the most enriched in immature neurons. **B.** Network analysis of the 9 transcripts highlighted in A with the most binding partners, blue indicates genes upregulated in neural progenitor cells and red indicates increased expression in immature neurons.

### Transcription factor networks and differential gene expression in neural progenitor cells and neurons

Based on the analysis described above, we were interested in determining whether differentially expressed transcription factors could operate as master regulators, controlling the differential expression of other transcripts in neural progenitor cells and immature neurons. We reasoned that master regulators might be identified based on 2 criteria: the number of their targeted genes and the number of interactions they have with other transcription factors, assuming that a transcription factor with more protein-protein interactions can indirectly regulate transcription of more genes. As a test case, we mined the ENCODE database and identified 126 transcription factors that could regulate the expression of our 635 differentially expressed transcripts (Figure 4A, Supplemental Data S4). Each transcription factor can regulate between 3 and 488 of the differentially expressed target genes, indicated as the size of the circle in Figure 4A. Three transcription factors, TAF1, SIN3A and MAX, regulate the most differentially expressed transcripts, with 488, 484, and 484 gene targets, respectively. We then conducted a protein-protein interaction network analysis, using STRING and Cytoscape. This revealed that only a subset of 4 of the 126 transcription factors is well connected in the network, indicated with hot colors in Figure 4A. EP300, with the highest network connectivity, has 79 interactions, followed by HDAC1 (73), MYC (67) and JUN (65). Indeed, these most highly connected transcription factors are recognized as master regulators, validating our strategy of using protein-protein interactions combined with the number of target genes to identify master regulators for cell proliferation and neuronal differentiation in our dataset of differentially expressed transcription factors.

**Figure 4.**
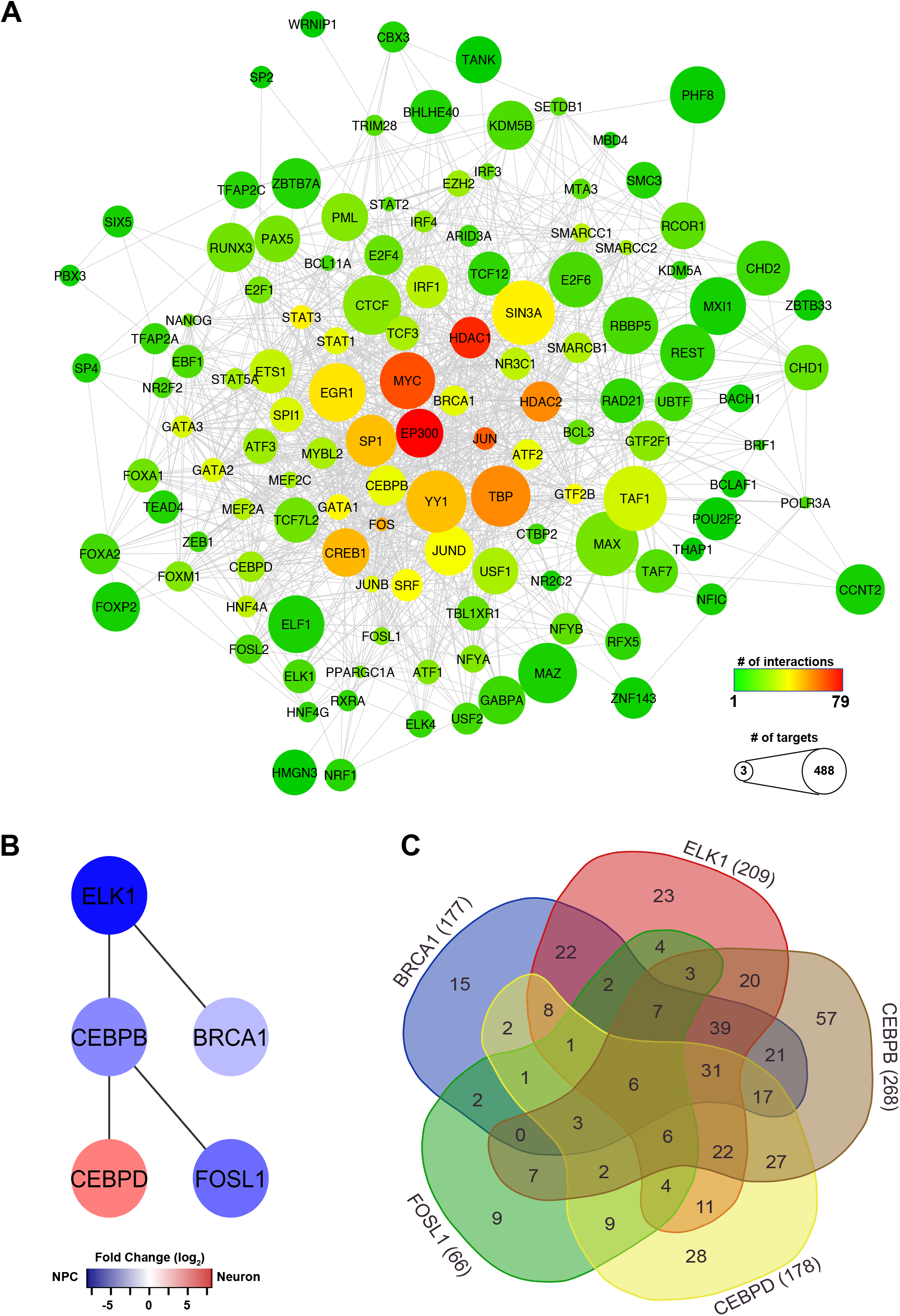
Transcription factor networks and differential gene expression in neural progenitor cells and neurons. **A.** Network analysis of the 127 differentially expressed transcription factors identified based on the ENCODE database. The size of the node reflects the number of differentially expressed transcripts that each transcription factor can potentially regulate. The color reflects the number of transcription factor binding partners. **B.** Network analysis of the most significantly differentially expressed transcription factors identified by the ENCODE database. Color refers to fold change. **C.** 5-way Venn diagram showing the numbers of overlapping transcripts targeted by the 5 differentially expressed transcription factors in B. For specific transcription targets, see Supplemental Table S2.

Using this same strategy on our dataset, we identified 6 differentially expressed transcription factors that themselves can regulate the other differentially expressed transcripts: BRCA1, ELK-1, CEBPB, CEBPD, FOSL1 and BRF1 (Figure 4B). CEBPD is enriched in immature neurons (Figure 4B, red), while the rest are enriched in neural progenitor cells (Figure 4B, blue). Using protein network analyses, we were surprised to find that 5 of these differentially expressed transcription factors, excluding BRF1 (transcription factor IIIB 90kDa subunit), interact with each other (Figure 4B). Their interactions suggest that the regulation of transcript expression is governed by a network of transcription factors, and these transcription factors might work together in a pathway or in synergy.

To further investigate whether there is cooperation among these 5 networked transcription factors with respect to their targeted genes, we generated a Venn diagram of the genes targeted by CEBPB (268), ELK-1 (209), CEBPD (178), BRCA1 (177) and FOSL1 (66) (Figure 4C, Supplemental Table S2). We found that 6 genes fell into the center overlap indicating they could be regulated by all 5 transcription factors. Four of these targets are enriched in immature neurons: APBA3 (fold change (FC) = 6.9), THUMPD (FC = 6.25), ELMOD3 (FC = 6.2), SLC39A3 (FC = 4.5), and the other 2 target genes are enriched in neural progenitor cells: C12orf57 (FC = −3.9) and MTMR4 (FC = −2.1). These 6 target genes, potentially regulated by 5 differentially expressed transcription factors, cover a diverse range of cellular processes from regulating neuronal health through transporting amyloid precursor proteins to regulating gene transcription through stabilizing tRNA. In particular, BRCA1 and ELK-1 together regulate a total of 270 target genes, 116 of which they both can act on (Figure 4C, blue and red regions, Supplemental Table S2). These data indicate that our network analysis approach identifies differentially-expressed potential master regulators, including both their protein interaction networks and transcriptional targets. We aim to examine the role of these master regulators, specifically BRCA1 and ELK-1, in neural progenitor cell fate in the developing brain.

### Visual experience alters BRCA1 and ELK-1 protein expression in neural progenitor cells

Of the networked transcription factors identified above, we chose to investigate further the roles of BRCA1 and ELK-1 which are enriched in neural progenitor cells (Figures 2C, 4B), interact with each other in a network of potential master regulators (Figure 4B), and regulate 270 of the differentially expressed targets in our dataset (Figure 4C), suggesting that they may play a role in neural progenitor cell fate determination. Because manipulating the dark/visual experience regime alters neural progenitor cell fate, we examined whether exposing animals to dark to enrich for neural progenitor cells affected protein expression of SOX2, BRCA1 and ELK-1. Western blots of the midbrain indicate that exposure to dark significantly increased SOX2 expression compared to levels seen in animals exposed to enhanced visual experience (Figure 5A), consistent with our previous findings that dark exposure increases SOX2+ neural progenitor cell proliferation (Bestman et al., 2012; Sharma and Cline, 2010). Similarly, BRCA1 and ELK-1 expression levels were significantly higher in animals exposed to dark compared to enhanced visual experience (Figure 5A), consistent with their higher transcript expression in neural progenitor cells (Figures 2C, 3B, 4B, Table 1).

**Figure 5.**
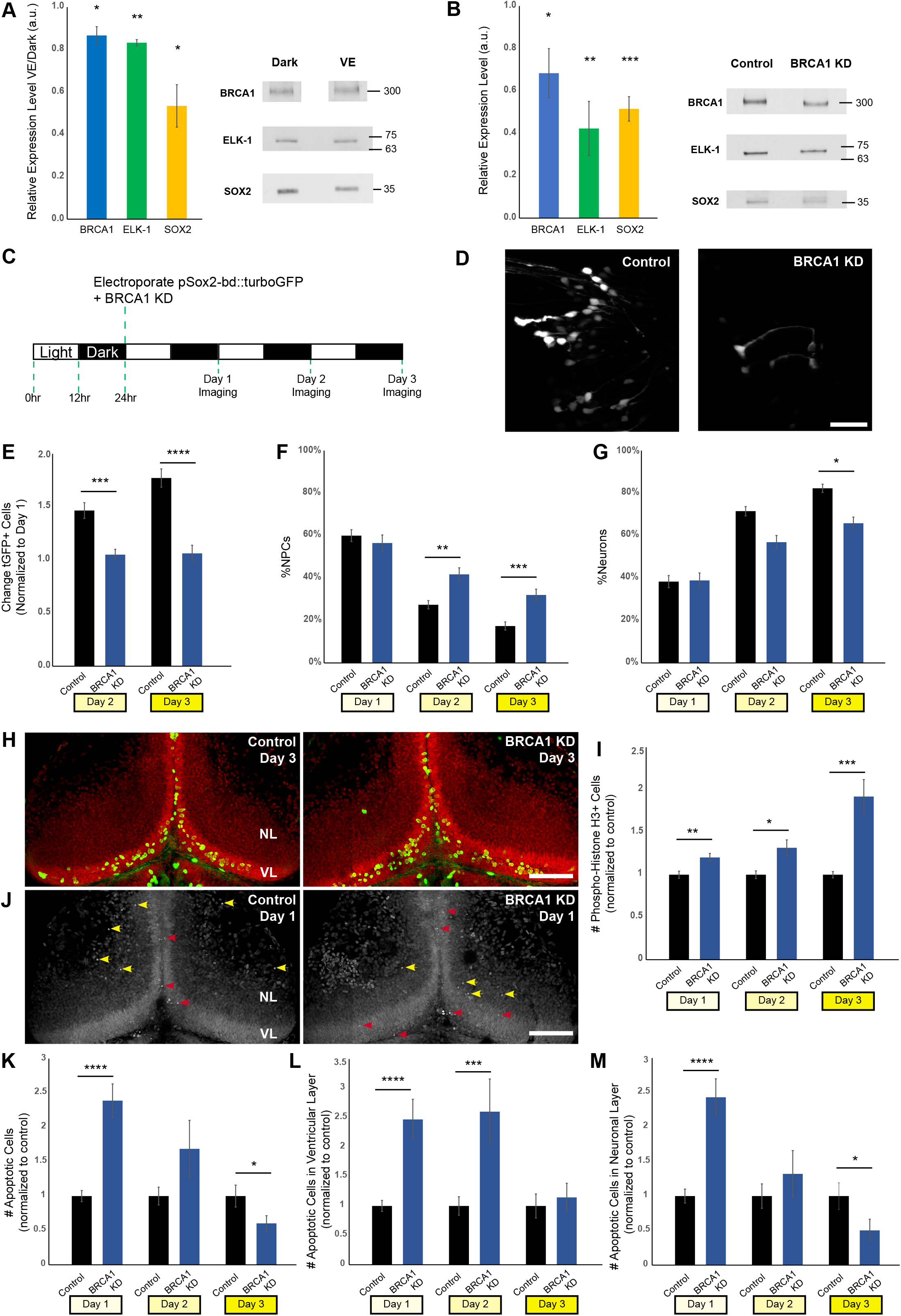
BRCA1 regulates the fate of neural progenitor cells. **A.** BRCA1 (blue), ELK-1 (green), and SOX2 (yellow) protein expression are increased in dark compared to enhanced visual stimulation. Quantitation (left) and representative western blots of BRCA1, ELK-1 and SOX2 in whole brain lysates (right). (n: BRCA1 (5); ELK-1 (4), SOX2 (4). **B.** BRCA1 knockdown lowers BRCA1 (blue, n=5), ELK-1 (green, n=6) and SOX2 (yellow, n=6) protein expression. **C.** Schematic diagram for *in vivo* imaging protocol for BRCA1 knockdown. After co-electroporation with tGFP and *brca1* morpholinos, animals were reared in a 12h light/12h dark cycle and imaged for 3 consecutive days by two-photon microscopy. **D.** Representative *in viv*o images of tGFP+ cells in the optic tectum of control and BRCA1 knockdown animals. Scale bar, 50um. **E.** BRCA1 knockdown significantly reduces the total number of tGFP+ cells compared to controls on both day 2 and 3. n=45-46 animals per condition. **F & G.** Of the cells shown in E, the percentage of neural progenitor cells significantly increased (**F**) and neurons decreased (**G**) with BRCA1 knockdown. **H.** Z-projection images showing pH3 immunolabelling (green) and SYTOX nuclear labelling (red) on day 3. Ventricular layer (VL) and neuronal layer (NL) are indicated. Scale bar: 100µm. **I.** BRCA1 knockdown significantly increased the total number of pH3+ cells in the tectum compared to controls over 3 days. n=37-48 animals per group/timepoint. **J.** Z-projection images showing SYTOX labelling to identify apoptotic cells on day 1. Red arrowheads indicate apoptotic neural progenitor cells in the ventricular layer (VL); yellow arrowheads specify apoptotic neurons in the neuronal layers (NL). Scale bar: 100µm. **K.** BRCA1 knockdown significantly affects the total number of apoptotic cells, increasing at day 1 and decreasing at day 3 as compared to controls. n=34-38 animals per group/timepoint. **L, M.** Of the cells shown in K, BRCA1 knockdown significantly increases apoptosis in neural progenitor cells on days 1 and 2 (**L**), and in neurons on day 1 (**M**). *p<0.05; **p<0.01; ***p<0.001; ****p<0.0001; 2-tail student t-test in A; 1-tail student t-test in B; Mann-Whitney U test in E, F, G, I, K, L, M.

### BRCA1 regulates the fate of neural progenitor cells

To examine the function of BRCA1 in neural progenitor cells, we used antisense morpholinos to knockdown BRCA1 by blocking the translation of *brca1* mRNA, thereby lowering BRCA1 protein expression, as confirmed by western blot (Figure 5B). In addition, BRCA1 knockdown (KD) decreased both ELK-1 and SOX2 protein levels (Figure 5B), suggesting that either BRCA1 directly regulates these proteins or that BRCA1 is required to maintain progenitor cell numbers. We then used *in vivo* time-lapse imaging to visualize the effect of BRCA1 KD on neural progenitor cells in optic tectum. Using the protocol illustrated in Figure 5C, tadpoles were reared under their normal 12h light/12h dark regime until stage 46, when their midbrains were electroporated with pSOX2-bd::turboGFP to express tGFP in SOX2+ neural progenitor cells. Simultaneously, midbrains were electroporated with either BRCA1 morpholino or control scrambled morpholino. We imaged tGFP+ neural progenitor cells and their neuronal progeny by collecting 2-photon images through the Z axis of the optic tectum at daily intervals for three days. We identified tGFP+ cells as neural progenitor cells by their soma positions lining the ventricle and their radial process tipped with an endfoot on the pial surface, and as tGFP+ neurons by their cell body shape and position in the neuronal cell body layer, and characteristic dendritic arbor morphology, as described (Bestman et al., 2012). We counted the total numbers of tGFP+ cells each day, and the numbers of neural progenitor cells and neurons (Figure 5D). BRCA1 KD blocked the normal increase in total number of tGFP+ cells seen under control conditions, maintaining a constant number of tGFP+ cells over the 3 days of imaging (Figure 5E), consistent with the decrease of the progenitor cell marker SOX2 seen by western blot (Figure 5B). We further examined whether BRCA1 KD alters the fate of neural progenitor cells, changing the cellular composition of tGFP+ progeny. BRCA1 KD increased the proportion of neural progenitor cells (Figure 5F) and decreased the proportion of neurons (Figure 5G) in the tGFP+ progeny at day 2 and day 3 compared to control. These data suggest that BRCA1 may affect both neural progenitor cell proliferation and differentiation of neural progenitor cells into neurons.

To examine whether BRCA1 KD blocked the increase in the total tGFP+ cells by blocking cell division, we immunolabeled cells for the cell division marker phospho-histone H3 (PH3) (Figure 5H, green cells). BRCA1 KD increased the number of PH3+ cells in the optic tectum compared to control morpholino conditions, indicating that BRCA1 KD did not block neural progenitor cell proliferation, but instead, increased the number of neural progenitor cells undergoing cell division (Figure 5I).

To investigate whether BRCA1 KD increases apoptosis as well as proliferation, thereby maintaining a constant total tGFP+ cell number over the 3 day period, we used SYTOX nuclear dye to identify apoptotic cells by condensed nuclear labelling (Figure 5J). BRCA1 KD resulted in a transient increase in apoptosis in the optic tectum compared to control animals (Figure 5K). To further dissect which cell type undergoes apoptosis, we categorized the apoptotic cells into neural progenitor cells and neurons based on their cell body localization. Compared to control, BRCA1 KD transiently increased apoptosis in neural progenitor cells at day 1 and day 2, and in neurons at day 1 (Figure 5L, 5M). These data show that BRCA1 KD increases the proliferative activity of neural progenitor cells and transiently increases the apoptotic activity of both neural progenitor cells and neurons. Together with the observations that BRCA1 KD blocks the normal increase in SOX2+ progeny, these data suggest that BRCA1 KD biases neural progenitor cells to undergo symmetric divisions, generating neural progenitor cell daughter cells, many of which become apoptotic along with some neurons. Together these data indicate that BRCA1 regulates the fate of neural progenitor cells.

### ELK-1 regulates the fate of neural progenitor cells

We examined the function of ELK-1 in neural progenitor cells using antisense morpholino to block the translation of *elk-1* mRNA, thereby decreasing ELK-1 protein expression, as shown by western blot (Figure 6A, green). ELK-1 KD also decreased levels of SOX2 protein similarly to BRCA1 (Figure 6A, yellow). Using a similar protocol as described in Figure 5C, we simultaneously electroporated the optic tectum with the SOX2bd::turboGFP plasmid and either *elk-1* morpholinos or control morpholinos. To assess the effect of ELK-1 knockdown (ELK-1 KD) on neurogenesis, we collected *in vivo* time-lapse images of tGFP+ neural progenitor cells and their neuronal progeny in the optic tectum (Figure 6B) at daily intervals for three days and analyzed changes in the numbers of tGFP+ neural progenitor cells and neurons as described above (Figure 5C). ELK-1 KD blocked the normal increase in tGFP+ cell numbers seen over 3 days *in vivo*, compared to control (Figure 6C, green bar). Furthermore, ELK-1 KD significantly increased the percentage of neural progenitor cells and decreased the percentage of neurons in tGFP+ progeny at both day 2 and day 3, compared to control (Figure 6D, 6E), indicating that ELK-1 KD alters the fate of neural progenitor cells.

**Figure 6.**
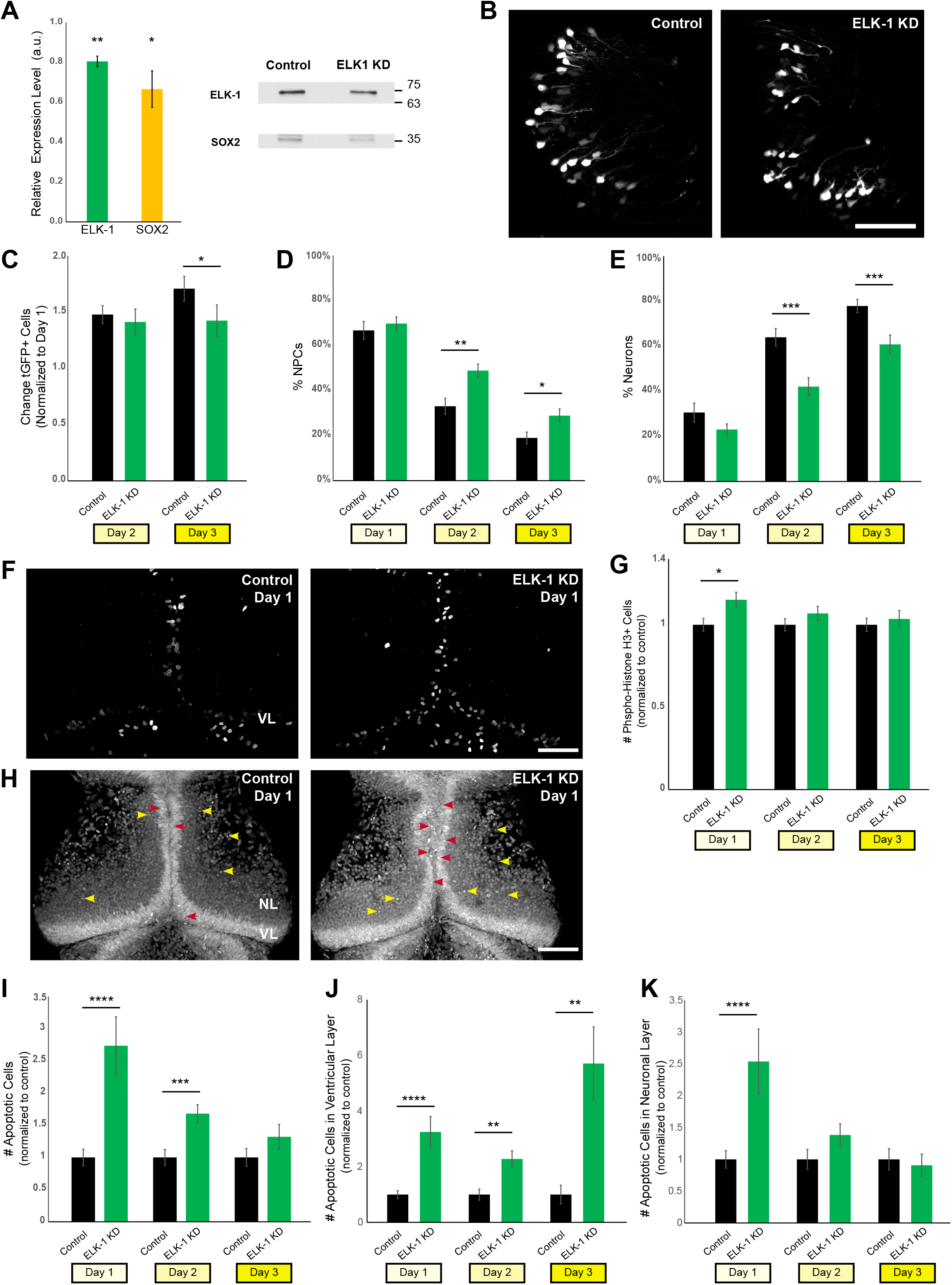
ELK-1 regulates the fate of neural progenitor cells. **A.** Knockdown with *elk-1* morpholino decreases ELK-1 protein expression (green, n=3) and reduces SOX2 levels (yellow, n=3). **B.** Representative *in vivo* images of tGFP+ cells in the optic tectum of control and ELK-1 knockdown animals. Experimental paradigm is the same as in Figure 5C. Scale bar: 100um**. C.** ELK-1 knockdown significantly reduces the total number of tGFP+ progeny number on day 3 compared to controls. n=12 animals per condition. **D & E.** Of the cells shown in C, ELK-1 knockdown significantly increases the percentage of neural progenitor cells (**D**) and decreases neurons (**E**). **F.** Confocal Z-projection of phospho-Histone H3 (pH3) immunolabelling in control and ELK-1 knockdown midbrains. Scale bar: 100µm. **G.** ELK-1 knockdown increases proliferative pH3-positive cells on day 1 compared to control. n=40-52 animals per group/timepoint. **H.** Z-projection of confocal microscopy images showing SYTOX nuclear labelling to identify apoptotic cells on day 1. Red arrowheads indicate apoptotic neural progenitor cells in the ventricular layer, yellow arrowheads signify apoptotic neurons. Scale bar: 100µm. **I.** ELK-1 knockdown significantly increases the total number of apoptotic cells on day 1 and 2 as compared to controls. n=39-51 animals per group/timepoint. **J, K.** Of the cells shown in I, ELK-1 knockdown significantly increased in apoptosis of neural progenitor cells across all days tested (**J**), and in neurons on day 1 (**K**) as compared to controls. *p<0.05; **p< 0.01; ***p<0.001; ****p<0.0001; 1-tail student t-test in A; Mann-Whitney U test in C, D, E, G, I, J, K.

To examine whether ELK-1 KD blocked the normal increase in tGFP+ cells over 3 days by blocking cell division, we counted PH3 immunolabeled cells in the optic tectum to identify dividing neural progenitor cells. ELK-1 KD transiently increased the proliferative activity at day 1, compared to control (Figure 6F, G). Furthermore, SYTOX labeling showed that ELK-1 KD increased apoptosis in the optic tectum (Figure 6H, I). Specifically, ELK-1 KD increased apoptosis in neural progenitor cells over all 3 days of the experiment (Figure 6J), and transiently increased apoptosis in neurons at day 1 compared to control (Figure 6K). These data show that ELK-1 KD increased the proportion of neural progenitor cells in the total tGFP+ pool. Underlying the increase in the proportion of neural progenitor cells was a transient increase in neural progenitor cell proliferation combined with increased apoptosis in neural progenitor cells over 3 days and a transient increase in apoptosis in neurons. Together, these studies suggest that ELK-1 KD drives neural progenitor cells to undergo symmetric divisions to expand the progenitor pool, while also increasing neural progenitor cell apoptosis. Similar to our findings with BRCA1 KD, our data indicate that ELK-1 regulates the fate of neural progenitor cells.

### Visual experience effects on neural progenitor cell fate depend on BRCA1

Our previous studies indicate that different visual experience conditions affect neural progenitor cell fate in the optic tectum, where exposure to dark increases progenitor cell proliferation and visual stimulation promotes neuronal differentiation. In the following experiments, we were interested in determining whether the effects of visual experience on neural progenitor cell fate require BRCA1 expression. Studies described above indicate that BRCA1 is more highly expressed in neural progenitor cells than neurons (Figure 4) and our *in vivo* imaging data suggest that BRCA1 governs neurogenesis by limiting neural progenitor cell proliferation and apoptosis in animals maintained in the normal 12h light/12h dark conditions. SOX2 and BRCA1 expression increase in the optic tectum in animals exposed to dark (Figure 5A), consistent with an increase in SOX2-, BRCA1-expressing neural progenitor cells.

To investigate whether different visual experience conditions require BRCA1 to affect neural progenitor cell fate, we monitored tGFP+ cells over three days *in vivo* using time-lapse imaging in animals that were maintained under control 12h light/12h dark conditions or maintained in the dark with or without BRCA1 knockdown. As schematized in the protocol in Figure 7A, stage 46 tadpole midbrains were electroporated simultaneously with pSOX2-bd::turboGFP to express tGFP in neural progenitor cells, and either *brca1* morpholinos or control scrambled morpholinos. We collected 2-photon images of tGFP+ neural progenitor cells and their neuronal progeny at daily intervals for three days. After baseline imaging on Day 1, animals were randomly divided into 2 groups, one of which was returned to control 12h light/12h dark conditions while the other group was maintained in the dark over the 3-day experiment except during imaging sessions. Representative images of tGFP+ cells imaged on day 3 are shown in Figure 7B. We counted the total number of tGFP+ cells each day and determined the proportion of neural progenitor cells and neurons based on cell morphology and position. We plotted the change in total tGFP+ cell numbers between Days 1 and 3, normalized to Day 1 (Figure 7C). First, we find that in animals exposed to the normal light/dark condition BRCA1 KD decreased tGFP+ cell numbers compared to controls (Figure 7C, gray vs. light blue bars), reproducing data shown in Figure 5E. Second, we found that BRCA1 KD blocked the dark-induced increase in tGFP+ cell numbers (Figure 7C, black vs. dark blue bars). These data show that the dark-induced increase in cell proliferation requires BRCA1. To further dissect how visual experience and BRCA1 protein expression are involved in regulating the fate of tGFP+ neural progenitor cells, we conducted a two-way ANOVA statistical analysis to test if there is an interaction between these two factors regarding tGFP+ cell numbers. The factorial experimental design we employed enabled us to perform such analysis. The profile plot shown in Figure 7D illustrates the relationship between the visual experience conditions and BRCA1 protein expression. Two-way ANOVA analysis reveals a statistically significant interaction between visual experience and BRCA1 protein expression on tGFP+ cell number. In control animals (Figure 7D, black line), the fold change in tGFP+ cells increased in the dark compared to 12h light/12h dark conditions. By contrast, when the BRCA1 KD animals were maintained in dark over the 3-day experiment (Figure 7D, blue line), tGFP+ cell numbers decreased compared to control animals. These data indicate that effects of visual experience on tGFP+ cell numbers depend on BRCA1 protein expression.

**Figure 7.**
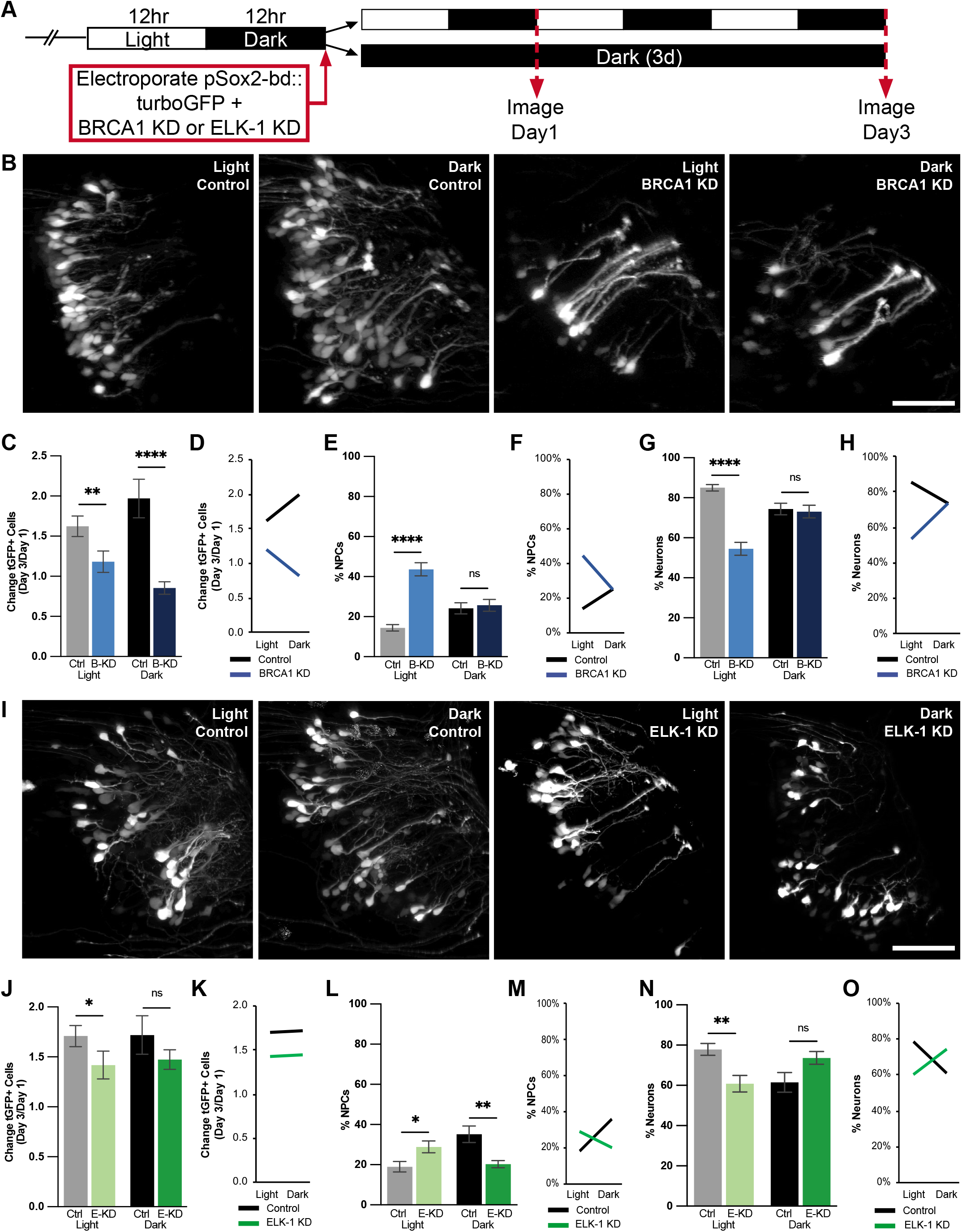
Visual experience effects on neural progenitor cell fate are mediated by BRCA1 and ELK-1. **A.** Schematic of treatment and imaging protocol. **B-H**. BRCA1 mediates the effects of visual stimulation on the fate of neural progenitor cells. **B.** Representative two-photon microscope *in vivo* images of tGFP+ cells in the optic tectum of animals in exposed to 12h light/12h dark conditions (light) or 3 days if continuous dark conditions (dark) with BRCA1 knockdown or animals treated with control morpholinos. Scale bar: 50um. **C-H**. Quantitative analysis of *in vivo* imaging data. **C.** BRCA1 knockdown (B-KD) blocked the normal increase in total GFP+ cells over the 3-day imaging period in animals exposed 12h light/12h dark conditions (light: grey bar vs light blue bar) and in animals exposed to dark (dark: black bar vs dark blue bar). n=17-24 animals per condition/timepoint. **D.** Profile plot of data in C demonstrating that the effect of visual experience on the total number of tGFP+ progeny is dependent on BRCA1. **E & G.** Of the cells shown in C, BRCA KD significantly increased the proportion of neural progenitor cells generated in animals exposed to 12h light/12h dark conditions (E. light: grey bar vs light blue bar) and decreased the proportion of neurons generated (G. light: : grey bar vs light blue bar). BRCA1 KD did not affect the proportion of neural progenitor cells (E) or neurons (G) in animals exposed to the dark condition (black vs dark blue bars). **F & H.** Profile plots of data in E and G demonstrating that the effect of visual experience on the fate of neural progenitor cells is dependent on BRCA1 protein expression. **I-O.** ELK-1 mediates the effects of visual stimulation on the fate of neural progenitor cells. **I.** Representative two-photon microscope *in vivo* images collected on day 3 of tGFP+ cells in the optic tectum of animals exposed to light or dark conditions with ELK-1 knockdown. Scale bar: 50um. **J-O.** Quantitative analysis of *in vivo* imaging data. **J.** ELK-1 knockdown (E-KD) blocks the normal increase in total GFP+ progeny in animals exposed to the exposed to 12h light/12h dark conditions (light: grey vs light green bars). ELK-1 knockdown in animals exposed to dark did not significantly affect total GFP+ progeny (black vs dark green bars). n=11-14 animals per condition/timepoint. **K.** Profile plot of factorial comparison of data in I, demonstrates that the effect of visual experience on the total number of tGFP+ cells is not dependent on ELK-1. **L.** Of the cells shown in J, ELK-1 KD significantly increased the proportion of neural progenitor cells under light conditions (grey vs light green bars) and significantly decreased neural progenitors under dark conditions (black vs dark green bars). **N**. Of the cells shown in J, ELK-1 knockdown significantly decreased the proportion of neurons under light conditions (grey vs light green bars) but did not significantly affect the proportion of neurons in animals maintained in the dark (black vs dark green bars). **M & O.** Profile plots of data in L and N demonstrate that the effect of visual experience conditions on the fate of neural progenitor cells and neurons depends on ELK-1 expression. *p<0.05, **p<0.01, ***p<0.001, ****p<0.0001; Mann-Whitney U test in C, E, G, J, L, N. Two-way ANOVA analysis was used in D, F, H, K, M, O.

We next examined whether the influence of visual experience on the proportion of neural progenitor cells is affected by BRCA1 expression. Data in Figure 5F show that under normal 12h light/12h dark conditions BRCA1 KD increases the proportion of neural progenitor cells in the tGFP+ cell population, and this result is independently reproduced in Figure 7E (gray vs. light blue bars). Visual experience with the 12h light/12h dark condition significantly increases neural progenitor cells when BRCA1 is knocked down, suggesting that BRCA1 normally limits the generation of neural progenitors under control visual experience conditions. In contrast, BRCA1 KD does not change the dark-induced proportion of NPCs (Figure 7E, black vs dark blue bars), suggesting that dark-induced proliferation is not sensitive to decreasing BRCA1 levels. Two-way ANOVA analysis shows a statistically significant interaction between visual experience and BRCA1 on the proportion of neural progenitor cells in the total tGFP+ cells (Figure 7F). The BRCA1 KD-facilitated increase in the proportion of neural progenitor cells only occurs in animals under normal 12h light/12h dark conditions while BRCA1 KD does not appear to effect the proportion of neural progenitor cells in animals maintained in the dark (Figure 7E, black vs dark blue bars, and Figure 7F). This suggests that BRCA1 may only be involved in neural progenitor cell fate in response to visual experience with light and might not have a role in increasing the proportion of neural progenitor cells when animals are maintained in the dark. Analysis of the proportion of neurons in the tGFP+ cell population suggests a similar conclusion. In animals under 12h light/12h dark conditions, BRCA1 KD significantly decreases the proportion of neurons in the tGFP+ cell population compared to controls (Figure 7G, gray vs. light blue bars) and there is a significant interaction between the visual experience conditions and BRCA1 protein expression on the proportion of neurons in the tGFP+ cells (Figure 7H), indicating that BRCA1 expression affects neuron number under normal light/dark conditions. However, as predicted from the neural progenitor cell data in Figure 7E, the proportions of neurons are similar in BRCA1 KD and control animals maintained in the dark (Figure 7G, black vs. dark blue bars), suggesting that BRCA1 protein expression is not involved in decreasing the proportion of tGFP+ neurons when animals are maintained in the dark. Together, these data indicate that the effects of visual experience on the fate of tGFP+ neural progenitor cells depend on BRCA1 protein expression.

### Visual experience effects on neural progenitor cell fate are mediated by ELK-1

To further understand the mechanisms by which visual experience conditions affect neural progenitor cell fate, we tested whether the effects of visual experience on neural progenitor fate are mediated by ELK-1. Thus far, our data indicate that ELK-1 is more highly expressed in neural progenitor cells than neurons (Figure 4), that exposing animals to dark conditions increases ELK-1 expression (Figure 5), and that ELK-1 limits neurogenesis by affecting both neural progenitor cell proliferation and apoptosis (Figure 6) in animals maintained in the normal 12h light/12h dark condition. Next, we used the protocol schematized in Figure 7A to determine whether the influence of visual experience conditions on cell proliferation and differentiation requires ELK-1 expression. Representative images of tGFP+ cells are shown for the control morpholino-treated animals and ELK-1 KD animals after 3 days of control or dark-exposed conditions (Figure 7I). Analysis of the imaging data show that ELK-1 KD decreased the number of tGFP+ cells in animals maintained in normal 12h light/12h dark conditions, independently reproducing the result shown in Figure 6, but ELK-1 KD did not significantly change tGFP+ cell numbers in animals exposed to dark (Figure 7J). Furthermore, two-way ANOVA statistical analysis of ELK-1 protein expression and visual experience conditions suggests that these two factors do not interact to regulate the number of tGFP+ cells, indicated by the two parallel lines in the profile plot (Figure 7K).

Recalling that maintaining animals in the dark increases neural progenitor cells (Figure 7E) and that ELK-1 KD increases the proportion of neural progenitor cells in animals under control conditions (Figure 6D), we next examined whether the effect of dark exposure on increasing neural progenitor cells depends on ELK-1 protein expression. Under normal visual experience conditions with 12h light/12h dark, ELK-1 KD increases the proportion of neural progenitor cells in the tGFP+ cell population compared to controls (gray vs. light green bars), independently replicating data shown in Figure 6D. These data indicate that ELK-1 normally limits the generation of neural progenitors under control visual experience conditions. Furthermore, ELK-1 KD blocked the dark-induced increase in the proportion of neural progenitor cells (Figure 7L, black vs. dark green bars), indicating that ELK-1 is required for the dark-induced expansion of the neural progenitor pool. We find a statistically significant interaction between the visual experience condition and ELK-1 protein expression on the proportion of neural progenitor cells in the tGFP+ cell population based on the two-way ANOVA analysis (Figure 7M). These data indicate that ELK-1 may be involved in neural progenitor cell fate in both visual experience with 12h light/12 h dark and continuous dark conditions. Analysis of the proportion of neurons in the tGFP+ cell population, indicates that in 12h light/12h dark conditions, ELK-1 KD significantly decreases the proportion of neurons compared to controls (Figure 7N, gray vs. light green bars) and there is a significant interaction between the visual experience condition and ELK-1 protein expression on the proportion of neurons in the tGFP+ cells (Figure 7O). However, ELK-1 KD did not significantly affect the proportion of neurons generated under dark conditions (Figure 7N, black vs dark green bars), indicating that ELK-1 is required specifically to mediate the effects of dark exposure on expanding the neural progenitor cell pool and the effects of light in promoting neuronal differentiation. Together, these data demonstrate that the effects of visual experience conditions on neural progenitor cell fate depends on ELK-1 protein expression.

## Discussion

This study profiled the transcriptomes of neuronal activity-induced proliferating neural progenitor cells and newly-differentiated immature neurons using RNA-seq in *Xenopus laevis* tadpole brain. Neuronal activity has been shown to regulate the fate of neural progenitor cells in the CNS in mammals and in non-mammalian vertebrates, such as frog tadpoles and fish (Bestman et al., 2015; Bestman et al., 2012; Hall and Tropepe, 2018; Madsen et al., 2000; Sharma and Cline, 2010; Sierra et al., 2015; Song et al., 2012). Here, we exposed tadpoles to dark or visual stimulation to bias the *in vivo* fate of SOX2+ neural progenitor cells in the optic tectum toward cell proliferation or differentiation, respectively, and used RNA-Seq of FAC sorted cells to determine the molecular signature of activity-dependent effects on neural progenitor cell fate. This experimental design enables us to detect differences in the transcriptomes of neural progenitor cells and immature neurons induced in the unperturbed neurogenic niche in the intact developing brain. Our transcriptome analysis identified 1,130 transcripts that were differentially expressed between neural progenitor cells and neurons. Data-mining and bioinformatic analyses revealed an overview of the potential roles of the differentially-expressed transcripts in neurogenesis. We identified a transcriptional network, including BRCA1 and ELK-1, that is comprised of differentially expressed transcripts. We propose that this differentially-expressed transcriptional network then cascades to regulate the expression of other differentially expressed transcripts in SOX2+ neural progenitor cells and their neuronal progeny during *in vivo* sensory experience-driven brain development. Using *in vivo* time-lapse 2-photon imaging, combined with translation-blocking antisense morpholinos, we showed that BRCA1 and ELK-1 affect neural progenitor cell fate, leading to overall decreased neurogenesis. Furthermore, by comparing the effect of exposure to dark or visual experience, we showed that BRCA1 and ELK-1 mediate sensory experience-dependent effects on neural progenitor cell fate. Together these studies provide a resource for transcriptomic profiles of enriched populations of neural progenitor cells and immature CNS neurons from the *X. laevis* midbrain, increase our understanding of cellular and molecular mechanisms underlying experience-dependent neural progenitor cell fate decisions in the *X. laevis* tadpole brain, and demonstrate roles for BRCA1 and ELK-1 in experience-dependent neurogenesis in the developing vertebrate brain.

### Isolation of cell populations enriched for Neural Progenitor Cells and neurons

Several studies have used genetically-expressed reporters to isolate and analyze neural progenitors from a variety of species, including Xenopus, Zebrafish and mice (Gaete et al., 2012; Kakebeen et al., 2020; Wang et al., 2011; Zupanc, 2021). Our strategy to isolate populations of neural progenitor cells and their neuronal progeny used a construct that requires binding of endogenous SOX2 to drive tGFP expression in SOX2+ neural progenitor cells. The HMG-Box transcription factor, SOX2, is expressed in neural progenitor cells in vertebrate brain, including Xenopus (Bestman et al., 2012; Gaete et al., 2012) and is necessary to maintain progenitor cell identity (Gotz et al., 2016). The fast maturation kinetics of tGFP and stability of the protein enabled us to label and isolate SOX2+ neural progenitor cells and their progeny within 24 hours after electroporation, and to image them *in vivo* over several days. Furthermore, we took advantage of prior studies showing that exposing tadpoles to dark or visual experience biased neural progenitor cell fate toward cell proliferation or differentiation, respectively (Bestman et al., 2012; Sharma and Cline, 2010). This allowed us to collect populations of FAC sorted tGFP+ cells that were enriched in neural progenitor cells or neurons, as validated by expression of canonical progenitor and neuronal markers (Figure 1, Table 1). Importantly, this strategy also allowed us to search for candidate mechanisms that might drive visual experience-dependent regulation of neural progenitor cell fates toward generating progenitor or neuronal progeny.

In the analysis shown in Figures 2-4, we mined through several databases in order to reveal the differences in transcriptomes between neural progenitor cells and immature neurons and the potential implications these differences represent. Recognizing that we are not analyzing pure populations of single neural cell types, our bioinformatic approaches emphasized network interactions, which weigh multiple components with known interactions or operating within known signaling pathways. Furthermore, our database analyses are built upon the abundant human data; since signaling pathways are largely conserved throughout evolution, the analyses based on human data are transferrable to *Xenopu*s and vice versa (Pires-daSilva and Sommer, 2003).

### Bioinformatic analysis identified functional categories and networks of differentially expressed genes

PANTHER’s functional categorization of the differentially expressed transcripts provided an overview of the differential expression patterns of these gene products and how they may play important roles in neurogenesis. Prominent categories of the differentially expressed transcripts include proteases and transcription factors. Differentially expressed transcripts in both of these categories include transcripts that have been shown in other systems to regulate progenitor cell fate and serve to validate our approach. The transcription factors we identified as enriched in neural progenitor cells, such as *e4f1*, *znf217* and *limx1b*, can promote proliferation or cell survival. For example, E4F1 and ZNF217 are reported to regulate cell survival and proliferation (Cowger et al., 2007; Paul et al., 2006). LMX1B, cooperatively with LMX1A, regulates proliferation and specification of midbrain dopaminergic (Marei et al., 2011). In contrast, the transcription factors we identified as enriched in immature neurons, such as *neuroD1*, *foxg1* and *evx1*, can induce the differentiation of neural progenitor cells into neurons. NeuroD1 induces terminal neuronal differentiation (Pataskar et al., 2015), while FOXG1 and EVX1 function in the specification of neuronal cell types (Thaeron et al., 2000; Yang et al., 2015).

Other identified transcripts are less well known with respect to progenitor cell fate regulation and may reveal additional molecular and cellular mechanisms involved in this context. For instance, 3 of the 15 differentially expressed proteases encode members of the ADAMTS (a disintegrin and metalloproteinase with thrombospondin motifs) family of extracellular proteases: *adamts-5*, *adamts-14* and *adamts-17*. *Adamts-17* is involved in maintaining survival and proliferation of cancer cells, suggesting a role for ADAMTS-17 in proliferation of neural progenitor cells, since many oncogenes first found in cancer also play a part during neurogenesis and vice versa (Jia et al., 2014). On the other hand, ADAMTS-5 enhanced neurite extension in immature neurons (Hamel et al., 2008). *Adamts-14*, a newer and less studied member of the ADAMTS family, may influence genetic predisposition for multiple sclerosis (Goertsches et al., 2005), but its potential role in neurogenesis is unknown.

Cell proliferation, cell differentiation and cell cycle are GO slim biological processes that were identified by PANTHER as prominent functional categories represented in the differentially expressed transcripts. These three processes are most relevant to maintaining neural progenitor cell self-renewability and neuronal differentiation. Among the differentially expressed transcripts in our dataset that are included in these processes, *brca1*, *tgfa* and *jak2*, which are enriched in neural progenitor cells, are known to promote cell proliferation (Kim et al., 2010; Prakash et al., 2015; Tropepe et al., 1997), while *sstr5*, which is decreased in neural progenitor cells, is known to inhibit cell proliferation (Barbieri et al., 2008). *pdgfa*, in addition to its known role promoting cell proliferation, also regulates the migration of newly differentiated cells (Nagel et al., 2004), consistent with our observations that *pdgfa* expression was increased in immature neurons. Together, this analysis identifies neurodevelopmental cellular processes that are associated with the differentially-expressed transcripts. Furthermore, while some of the differentially expressed transcripts in these functional protein categories are known players in cell proliferation and neuronal differentiation, essentially validating our strategy, this analysis also supports hypotheses implicating candidates that are less-well known in neurodevelopmental processes.

### Protein-Protein interaction networks identify key players in neurodevelopment among the differentially expressed transcripts

In an effort to identify differentially expressed transcripts that may be important for neural progenitor cell fate and neuronal differentiation, we applied a strategy to identify the encoded proteins which have the most protein-protein interactions, based on the likelihood that these predicted protein interaction networks would play a regulatory role in the fates of neural progenitor cells and neurons. This analysis identified nine differentially expressed transcripts whose protein products had more than 20 interactions: ACTA2, BMP4, JAK2 and BRCA1, which were enriched in neural progenitor cells, and ITGA2, VEGFA, FGF2, AURKB and NFKB1, which were enriched in immature neurons. It is interesting to note that these highly connected hub proteins also interact with each other, supporting the idea of network-based regulation of fundamental neurodevelopment events such as cell fate.

These hub proteins are embedded in networks with key roles in neurodevelopment. ITGA2 (Integrin*α*2) has 40 protein interaction partners among the network of 458 proteins generated from our differentially expressed transcripts. Integrin*α*2 facilitates migration of differentiating embryonic stem cells and iPSCs (induced pluripotent stem cells) by remodeling extracellular matrix and initiating intracellular signaling cascades (Li et al., 2011). In addition to ITGA2, FGF2 (fibroblast growth factor 2) and VEGFa (vascular endothelial growth factor A), with 23 and 24 interactions in the network, respectively, are growth factors that regulate neuronal differentiation (Cavanagh et al., 1997; Louvi and Artavanis-Tsakonas, 2006; Rosenstein et al., 2003). VEGFa may also be involved in migration of newly differentiated neurons (Wang et al., 2015). We have previously demonstrated a role for *fgf2* in neurogenesis in *Xenopus* optic tectum, where FGF2 knockdown increased cell proliferation (Bestman et al., 2015), indicating that FGF2 maintains progenitor cell capacity for self-renewal (Sanalkumar et al., 2010). NF-kB1 (nuclear factor of kappa light polypeptide gene enhancer in B-cells 1) regulates neuronal differentiation in the adult mouse hippocampus and maintains cell survival (Denis-Donini et al., 2008). It is interesting to note that NF-kB1 signaling activity is regulated by Aurkb (Aurora kinase B) (He et al., 2015), another highly interconnected protein whose transcript is enriched in the immature neurons. In addition to these 5 transcripts that are enriched in immature neurons in our dataset, *brca1* (breast cancer 1) and *jak2* (Janus kinase 2, a non-receptor tyrosine kinase) are enriched in neural progenitor cells, and their proteins each had 28 and 29 interaction partners in this network, respectively (Figure 3B). BRCA1 can regulate the expression and modulate the activity of JAK2 (Gao et al., 2001). Following the BRCA1 signaling pathway, JAK2 activates the STAT signaling cascade, and *stat2* itself is enriched in neural progenitor cells. The JAK/STAT signaling cascade modulates proliferation of neural progenitor cells (Chung et al., 2013; Kang and Kang, 2008). *bmp4* (bone morphogenetic protein 4), also enriched in neural progenitor cells, has 25 interactions in this network and has been reported to maintain self-renewal of mouse embryonic stem cells (Zhang et al., 2013).

Our analysis of the protein-protein interaction network indicates that proteins generated from the differentially expressed transcripts occupy important positions within network nodes regulating cell proliferation, differentiation and survival, with distinct functions in regulating neurogenesis. For instance, several proteins in this network maintain progenitor self-renewal by promoting cell proliferation, suppressing neuronal differentiation, or enhancing neural progenitor cell survival. Other proteins in the network induce neuronal differentiation by increasing pro-neural gene expression or facilitating cell migration. It is interesting to note that proteins generated from differentially expressed transcripts are themselves well-connected in a network of other proteins derived from differentially expressed transcripts, consistent with coordinated transcriptional control generating functional protein interaction networks.

### Transcription factor networks that mediate activity-dependent control of neural progenitor cell fate

A major interest in our bioinformatic analysis is to identify differentially expressed master transcriptional regulators that could function in a network to regulate many of the other transcripts that were differentially expressed between neural progenitor cells and immature neurons in response to different visual stimulation conditions. As a proof-of-principle analysis, we first identified the transcription factors that are known to regulate the differentially expressed transcripts in our dataset. We then showed that using dual criteria to search for transcription factor master regulators based on 1. the capacity to regulate the majority of the differentially expressed transcripts in our dataset and 2. the number of interaction partners, successfully identified well-known transcriptional master regulators, including a network of highly interconnected transcriptional regulators composed of EP300, HDAC1, MYC, and JUN. Indeed, each of these transcriptional regulators has been shown to play roles in cell proliferation, differentiation, and cell fate determination. EP300 (E1A binding protein p300) acts as both a transcriptional co-activator and a histone acetyltransferase, positively regulating histone acetylation to initiate transcription. EP300 regulates transcription of both pluripotency genes (*c-myc, c-myb, creb, c-jun, and c-fos*) and neural genes (*pax6*, *sox1*, *zic2*, and *znf521*) (Chen et al., 2011; Chen et al., 2010; Qiao et al., 2015), indicating that EP300 can affect both cell proliferation and neural differentiation. HDAC1 (histone deacetylase 1) is a transcriptional regulator that epigenetically represses gene transcription. Inhibition of HDAC activity can both maintain pluripotency of human embryonic stem cells and inhibit neural differentiation, depending on its targets (Qiao et al., 2015). HDAC1 indirectly increases in c-Myc protein levels (Luo et al., 2000), which then increases self-renewability of neural progenitor cells (Zheng et al., 2008). c-Myc is thought to regulate expression of 15% of all genes (Gearhart et al., 2007). Both c-Myc and Jun regulate cell proliferation by regulating expression of Cyclins, and over-expression of c-Jun represses p53 expression, enhancing cell proliferation (Schreiber et al., 1999). These 4 most-connected transcription factors from our network of 126 transcription factors are recognized as master regulators of various cellular processes during development, validating our approach using protein-protein interactions and the number of target genes to identify master regulators in cell proliferation and neuronal differentiation.

We then applied this dual criteria strategy to our set of differentially-expressed transcription factors and identified five candidate master regulators of neural progenitor cell proliferation and differentiation that each have multiple interaction partners and together form a network: CEBPD, FOSL1, CEBPB, ELK-1, and BRCA1. Are these likely candidates to mediate sensory experience-dependent effects on transcript expression? Activity-dependent regulation of cell proliferation and neuronal differentiation are mediated by intercellular signaling between neurons and neural progenitor cells that initiates ERK/MAPK signaling (Kirischuk et al., 2017; Ma et al., 2009). ERK/MAPK signaling induces transcription of immediate early genes in neural progenitor cells, which in turn can induce expression of diverse genes (Yap and Greenberg, 2018). It appears that this network is well positioned to transduce activity-triggered intercellular signals to affect neural progenitor cell fate. Four members of the network, FOSL1, CEBPB, ELK-1 and BRCA1, are enriched in neural progenitor cells and interact with each other. FOSL1 (fos-like antigen 1) binds c-Jun to form the activator protein 1 (AP-1) transcription complex and promotes cell cycle progression (Hess et al., 2004). Although FOSL1 has not yet been shown to regulate proliferation of neural progenitor cells, it does regulate cancer cell proliferation and metastasis (Vallejo et al., 2017) (Pennanen et al., 2011). CEBPB (CCAAT/enhancer binding protein (C/EBP), beta) promotes self-renewal and proliferation of neural progenitor cells, survival of new-born neurons, and can bias the fate of cortical progenitor cells toward neurons (Pulido-Salgado et al., 2015). CEBPB also works in synergy with ELK-1 by interacting with serum response factor to transactivate *c-fos* (Hanlon et al., 2000).

BRCA1 is a multifunctional protein, widely known as a tumor suppressor, with roles in genome stability, checkpoint control, replication fork stability and DNA double strand break repair via homologous recombination (Densham and Morris, 2017; Frappart et al., 2007; Prakash et al., 2015). In mice, BRCA1 null is embryonic lethal and results in developmental deficits before neural tube closure, consistent with impaired repair of double stranded DNA breaks during rapid cell division in the embryo (Gowen et al., 1996; Orii et al., 2006). Spatial and temporal control of BRCA1 loss of function in mice demonstrated that animals with deficient machinery for homologous recombination mediated DNA repair have neurodevelopmental defects, and specifically that BRCA1 plays a role in neurogenesis (Pao et al., 2014; Pulvers and Huttner, 2009). BRCA1’s function repairing double stranded DNA damage is thought to be particularly important in neural progenitor cells because their high rate of proliferation makes them prone to DNA double strand breaks (Orii et al., 2006). In addition to its role in DNA repair, BRCA1 is a component of core transcriptional machinery, where it can act as a transcriptional activator or repressor, depending on its interaction partners (Mullan et al., 2006). BRCA1 is an upstream regulator of *elk-1,* and ELK-1 interacts with BRCA1 to augment BRCA1’s growth suppressive function in cancer cells (Maniccia et al., 2009) (Chai et al., 2001). In addition, BRCA1 can regulate the expression and modulate the activity of JAK2 (Gao et al., 2001), which we identified as a differentially expressed transcript with one of the highest number of protein-protein interactions, and can reportedly induce cell proliferation by activating promoters of *c-fos* and *c-myc*, an IEG, which itself was differentially expressed and has a high number of protein-protein interactions.

ELK-1 is expressed in SOX2+ neural progenitor cells and neurons throughout development and in adult animals (Wells et al., 2011). ELK-1 is phosphorylated by MAPKs, including ERK, resulting in translocation of pELK-1 to the nucleus and induced transcription of diverse target genes. The specificity of ELK-1-regulated transcriptional responses is likely due to recruitment of specific co-activators, such as CREB binding protein, p300, and serum response factor, regulating pluripotency, apoptosis, proliferation and survival in neural progenitors and synaptic plasticity in neurons (Besnard et al., 2011). Nuclear translocation of pELK-1 in SOX2+ neural progenitors induced transcription of IEGs, including *egr1* (aka *zif266*) and *c-fos*, as well as other targets pertaining to proliferation and pluripotency, such as *sox2, oct4, nanog* (Wells et al., 2011). This convergence of extracellular signaling to IEGs and *sox2* suggests a mechanism for extracellular activating signals to regulate neural progenitor cell fate.

CEBPD, (CCAAT/enhancer binding protein (C/EBP), delta) has been shown to inhibit cell proliferation by down-regulating c-Myc and cyclins and increasing expression of differentiation-related genes (Pulido-Salgado et al., 2015). This suggests that CEBPD may inhibit cell proliferation and drive neuronal differentiation, consistent with its increased expression we observed in immature neurons. In combination with the enriched expression of neuroD1 and fgf2, that are known to induce terminal neuronal differentiation (Louvi and Artavanis-Tsakonas, 2006), increased expression of CEBPD in response to visual experience in tadpoles may promote neural progenitor cells to exit the cell cycle and differentiate into neurons.

In summary, these 5 differentially expressed transcriptional regulators affect cell proliferation, differentiation, and survival of neural progenitor cells or immature neurons, likely functioning as a network to regulate many of the differentially expressed transcripts in our dataset. Together these data suggest that our transcriptome analysis identified factors and mechanisms that may regulate neural progenitor cell proliferation and neuronal differentiation in the developing brain in response to activity-driven cues.

### BRCA1 and ELK-1 knockdown occlude the effects of visual experience on neural progenitor cell fate

We identified *brca1* and *elk-1* as transcripts with greater expression in neural progenitor cells compared to neurons. Protein expression of BRCA1 and ELK-1 is greater in animals exposed to dark compared to those exposed to visual stimulation, consistent with greater transcript expression in progenitors. The increased BRCA1 and ELK-1 in western blots is also consistent with an expansion of the progenitor pool, shown in in Figures 6 and 7, and in our previous studies (Bestman et al., 2012; Sharma and Cline, 2010). In addition, our bioinformatic analysis indicated that BRCA1 and ELK-1 are members of a network of differentially expressed transcriptional regulators which may in turn target a large proportion of the transcripts that are differentially expressed between neural progenitors and neurons in our study, suggesting that visual experience conditions lead to sequentially amplifying effects on differential transcript expression in neural progenitors and neurons. Following these observations, our *in vivo* time-lapse imaging data showed that both BRCA1 and ELK-1 knockdown led to a net decrease in neurogenesis by transiently increasing apoptosis of both neural progenitor cells and neurons. Finally, we find that effects of visual experience conditions, both light and dark, on cell numbers, expansion of the progenitor pool and neuronal differentiation require BRCA1 and ELK-1.

We targeted BRCA1 knockdown specifically to the optic tectum of stage 46 tadpoles to avoid early lethality and large-scale neurodevelopmental defects reported in other studies (Gowen et al., 1996; Orii et al., 2006; Pao et al., 2014; Pulvers and Huttner, 2009) and to assess direct effects of BRCA1 manipulation in animals under different visual experience conditions. The overall decrease in tGFP+ cell numbers with BRCA1 KD seen with *in vivo* imaging follows the rapid increase in apoptosis, which was detected principally in neural progenitor cells, consistent with BRCA1’s role in homologous recombination-mediated DNA repair in proliferative cells in rodent brain (Orii et al., 2006; Pao et al., 2014; Pulvers and Huttner, 2009). The increased apoptosis seen with BRCA1 KD results in a change in the proportion of neural progenitor cells and neurons detected in the *in vivo* imaging data. In control animals, the proportion of neural progenitor cells decreases over 3 days of imaging, as the proportion of neurons increases reciprocally, but apoptosis in neural progenitor cells blocked the normal increase in neuronal differentiation, reducing the proportion of neurons in the tGFP+ population and paradoxically increasing the proportion of neural progenitors. It is interesting that we observe an increase in pH3 labeling in response to BRCA KD, suggesting that BRCA1 KD does not interfere with S-phase DNA replication.

ELK-1 KD decreased SOX2 in western blots from tectum and increased neural progenitor cell apoptosis, consistent with the observation that ELK-1 KD decreases expression of ‘stemness’ genes, such as *sox2, oct4, nanog* (Sogut et al., 2021). These studies suggest that ELK-1 KD drives neural progenitor cells to expand the progenitor pool through symmetric divisions, while also increasing neural progenitor cell apoptosis. BRCA1 and ELK-1 interact and this interaction enhances BRCA1 function, suggesting that ELK-1 may function downstream of BRCA1 (Chai et al., 2001). Our observation that BRCA1 KD decreases ELK-1 expression is consistent with known transcriptional regulation of ELK-1 by BRCA1 (Maniccia et al., 2009).

Comparing the effect of BRCA1 or ELK-1 knockdown under light or dark conditions indicates that the visual experience-dependent effects on neural progenitor fate are mediated by BRCA1 and ELK-1. In experiments with BRCA1 KD compared to control, the effects of visual experience on tGFP+ cell numbers are opposite in the two conditions. Under control conditions tGFP+ cell numbers increase over the 3-day exposure to dark, but with BRCA1 KD, tGFP+ cell numbers decrease after dark exposure. These data indicate that the visual experience-dependent increase in cell numbers requires BRCA1. When we analyzed the effect of visual experience on the proportion of neural progenitor cells and neurons, BRCA1 KD increased the proportion of neural progenitor cells and decreased the proportion of neurons in the tGFP+ population in response to visual stimulation provided with the 12h light/12h dark condition. This altered cell fate suggests that BRCA1 is required for the visual stimulation induced differentiation of neurons, such that with decreased BRCA1, neural progenitor cells fail to differentiate and their relative numbers increase. Similarly, we find that the influence of visual stimulation conditions on the proportions of neural progenitor cells and neurons is significantly altered by ELK-1 KD, indicating the ELK-1 is required for both the increased neural progenitor cell proliferation in the dark and the visual stimulation-induced neuronal differentiation. These effects of ELK-1 on proliferation and differentiation may be mediated by different ELK-1 targets and signaling pathways. Together these data indicate that BRCA1 and ELK-1 regulate neural progenitor cell fate, through both shared and diverse molecular and cellular pathways.

The interplay between BRCA1 and ELK-1 is particularly interesting in light of their shared roles in neural progenitor cell fate in response to visual activity. Animals exposed to visual stimulation have decreased expression of BRCA1, ELK-1 and SOX2 compared to animals exposed to dark. Furthermore, BRCA1 KD decreases ELK-1 expression, consistent with other studies indicating that BRCA1 negatively regulates *elk-1* transcription (Maniccia et al., 2009). In addition, BRCA1 and ELK-1 interact and this interaction enhances BRCA1 function, suggesting that ELK-1 may function downstream of BRCA1 (Chai et al., 2001). Decreased levels of BRCA1 and ELK-1 in the tectum increased neural progenitor cell apoptosis, consistent with the observation that ELK-1 KD decreases expression of ‘stemness’ genes, such as *sox2, oct4, nanog* (Sogut et al., 2021). These studies suggest that BRCA1 KD and ELK-1 KD drive neural progenitor cells to expand the progenitor pool through symmetric divisions, while also increasing neural progenitor cell apoptosis.

Together, BRCA1 and ELK-1 can potentially regulate 270 differentially expressed transcripts in our dataset, with 116 shared targets. Furthermore, BRCA and ELK-1 have the capacity to operate in a network with the 3 other differentially expressed transcriptional regulators, CEBPB, CEBPD, and FOSL1, to target a total of 409 transcripts. Of these, 6 transcripts can be regulated by all 5 of the differentially-expressed networked master regulator transcription factors. These 6 target genes, *apab3, thumpd, elmod3, slc39a2, c12orf57,* and *mtmr4,* are relatively less known and their functions range across diverse cellular processes, including cytoskeletal regulation, apoptosis, TGF*β* signaling and transcriptional regulation. Since little is known about these six genes in neuronal development, their potential involvement in BRCA1-and ELK-1-mediated regulation of neural cell fate remains to be discovered.

In summary, we used visual experience to manipulate neural progenitor cell fate and profiled the transcriptomes of the resulting neural progenitor cells and immature neurons. We identified a set of 1,130 differentially expressed transcripts, including a network of 5 master regulator transcription factors. Of these, we demonstrate that BRCA1 and ELK-1 play key roles in regulating neurogenesis in response to sensory input in the developing tadpole brain.

## Acknowledgements

Supported by TSRI CIRM Training Grant #01165 (L-C.H)., Dart Neuroscience, the NIH (NS076006, NS114975, EY EY011261, EY027437, EY031597) and an endowment from the Hahn Foundation to H.T.C. We thank members of the Cline lab for helpful comments and discussion, and both the Fluorescence Activated Cell Sorting Core and the Next Generation Sequencing Core at The Scripps Research Institute.

## Author contributions

L-CH, HTC designed experiments; L-CH, HH, ACT conducted experiments and data analysis; HTC supervised research; L-CH, CRM, HTC prepared figures and wrote the paper. All authors reviewed the final manuscript.

## Supplemental Data

**Supp figure 1.**
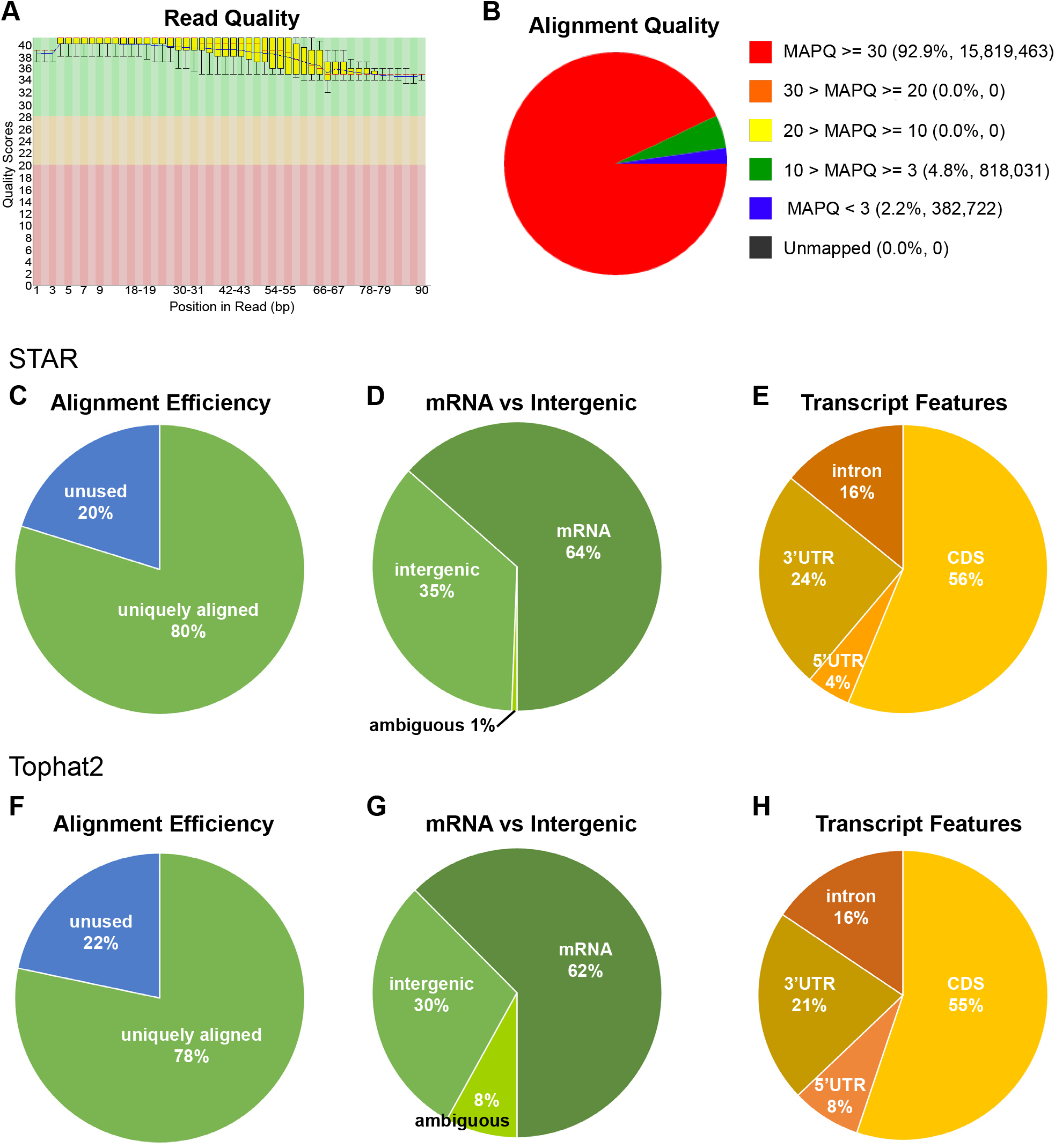
Read quality and alignment of differential expression dataset. **A.** Read quality of entire dataset including 3 biological replicates of both NPCs and immature neurons. **B.** MAPQ alignment quality using SAMStat. **C-E.** Details of alignment of reads against *Xenopus laevis* genome, using STAR. **C.** 80% of RNA-seq reads in average are uniquely aligned to the genome scaffolds, indicating the specificity of the reads to *Xenopus laevis* genome. **D.** Against genome scaffold (J-strain v9.1) and gff3 file, 64% of the aligned reads in average belong to mRNA; while 35% of the aligned reads in average belong to the regions between mRNA, ie. intergenic region. **E.** Percentage of reads aligned to the features in the transcripts. **F-H.** Details of alignment of reads against *Xenopus laevis* genome, using TopHat2. **F.** 78% of RNA-seq reads in average are uniquely aligned to the genome scaffolds, indicating the specificity of the reads to *Xenopus laevis* genome. **G.** Against genome scaffold (J-strain v9.1) and gff3 file, 62% of the aligned reads in average belong to mRNA; while 30% of the aligned reads in average belong to the regions between mRNA, ie. intergenic region. **H.** Percentage of reads aligned to the features in the transcripts.

**Supp figure 2.**
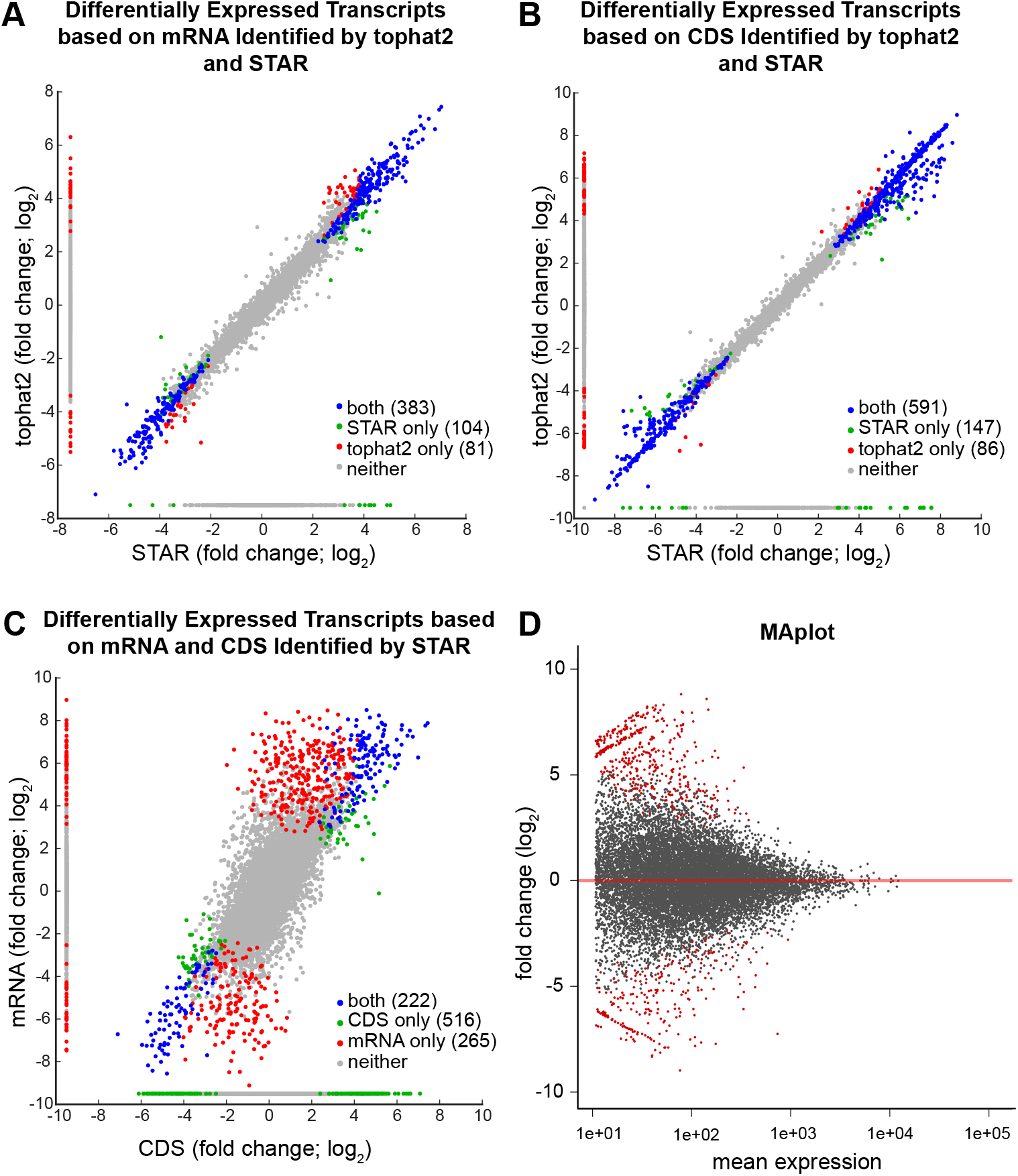
Comparison of the differentially expressed transcripts between neural progenitor cells and immature neurons, identified using STAR and TopHat2. **A.** A scatter plot showing the correlation of the fold change of transcript expression in immature neurons in comparison to neural progenitor cells based on mRNA between STAR and TopHat2. **B.** A scatter plot showing the correlation of the fold change of transcript expression in immature neurons in comparison to neural progenitor cells based on coding region (CDS) between STAR and TopHat2. **C.** A scatter plot showing correlation of the fold change of transcript expression in immature neurons in comparison to neural progenitor cells based on mRNA and CDS, using STAR. **E.** MA plot shows the mean expression of transcripts vs its fold change (log2) between neural progenitor cells and immature neurons. The differentially expressed genes are indicated in red with adjusted p-value < 0.1.

**Supplemental Table S1.** Read alignment against *Xenopus laevis* genome, using TopHat2 and STAR (**refers to methods)**

| **tophat 2** | **neural progenitor cells** |  |  | **immature neurons** |  |  |
| --- | --- | --- | --- | --- | --- | --- |
|  | **sample 1** | **sample 2** | **sample 3** | **sample 1** | **sample 2** | **sample 3** |
| **raw reads** | 17,590,066 | 19,990,928 | 16,804,899 | 17,523,604 | 19,423,625 | 19,859,802 |
| **trimmed reads** | 14,464,497 | 16,044,500 | 14,953,754 | 15,741,735 | 17,349,543 | 17,800,922 |
| **uniquely mapped reads** | 11,444,705 | 13,710,391 | 11,517,049 | 12,503,563 | 13,537,406 | 13,777,726 |
| **reads with MAPQ >= 30** | 10,477,365 | 12,427,637 | 10,572,169 | 11,509,693 | 12,276,215 | 12,545,706 |
| **reads counted for mRNA** | 6,947,287 | 8,048,157 | 7,111,316 | 7,779,759 | 8,132,974 | 8,324,947 |
| **reads counted for intergenic region** | 3,637,589 | 4,467,038 | 3,520,628 | 3,829,538 | 4,244,592 | 4,307,845 |
| **reads counted for CDS** | 3,118,975 | 4,485,633 | 4,392,286 | 3,628,992 | 4,439,246 | 5,125,655 |
| **reads counted for 3UTR** | 1,257,093 | 1,807,527 | 1,660,929 | 1,446,179 | 1,774,019 | 1,865,073 |
| **reads counted for 5UTR** | 589,252 | 843,930 | 590,526 | 676,117 | 779,413 | 668,988 |
| **reads counted for intron** | 1,911,967 | 911,067 | 467,575 | 2,028,471 | 1,140,296 | 665,231 |

| **STAR** | **neural progenitor cells** |  |  | **immature neurons** |  |  |
| --- | --- | --- | --- | --- | --- | --- |
|  | **sample 1** | **sample 2** | **sample 3** | **sample 1** | **sample 2** | **sample 3** |
| **raw reads** | 17,590,066 | 19,990,928 | 16,804,899 | 17,523,604 | 19,423,625 | 19,859,802 |
| **uniquely mapped reads** | 13,273,346 | 15,819,463 | 13,406,592 | 14,520,781 | 15,568,590 | 15,937,605 |
| **reads counted for mRNA** | 8,422,751 | 9,655,249 | 8,480,161 | 9,388,103 | 9,747,548 | 9,914,191 |
| **reads counted for intergenic region** | 4,767,796 | 6,071,063 | 4,853,417 | 5,045,312 | 5,728,889 | 5,935,165 |
| **reads counted for CDS** | 3,796,644 | 5,316,712 | 5,172,041 | 4,304,968 | 5,259,977 | 6,033,870 |
| **reads counted for 3UTR** | 1,561,072 | 2,245,761 | 2,060,704 | 1,792,487 | 2,201,242 | 2,312,324 |
| **reads counted for 5UTR** | 757,445 | 1,084,443 | 764,621 | 868,924 | 1,001,476 | 862,520 |
| **reads counted for intron** | 2,307,590 | 1,008,330 | 482,795 | 2,421,724 | 1,284,853 | 705,477 |

**Supplemental Table S2.**
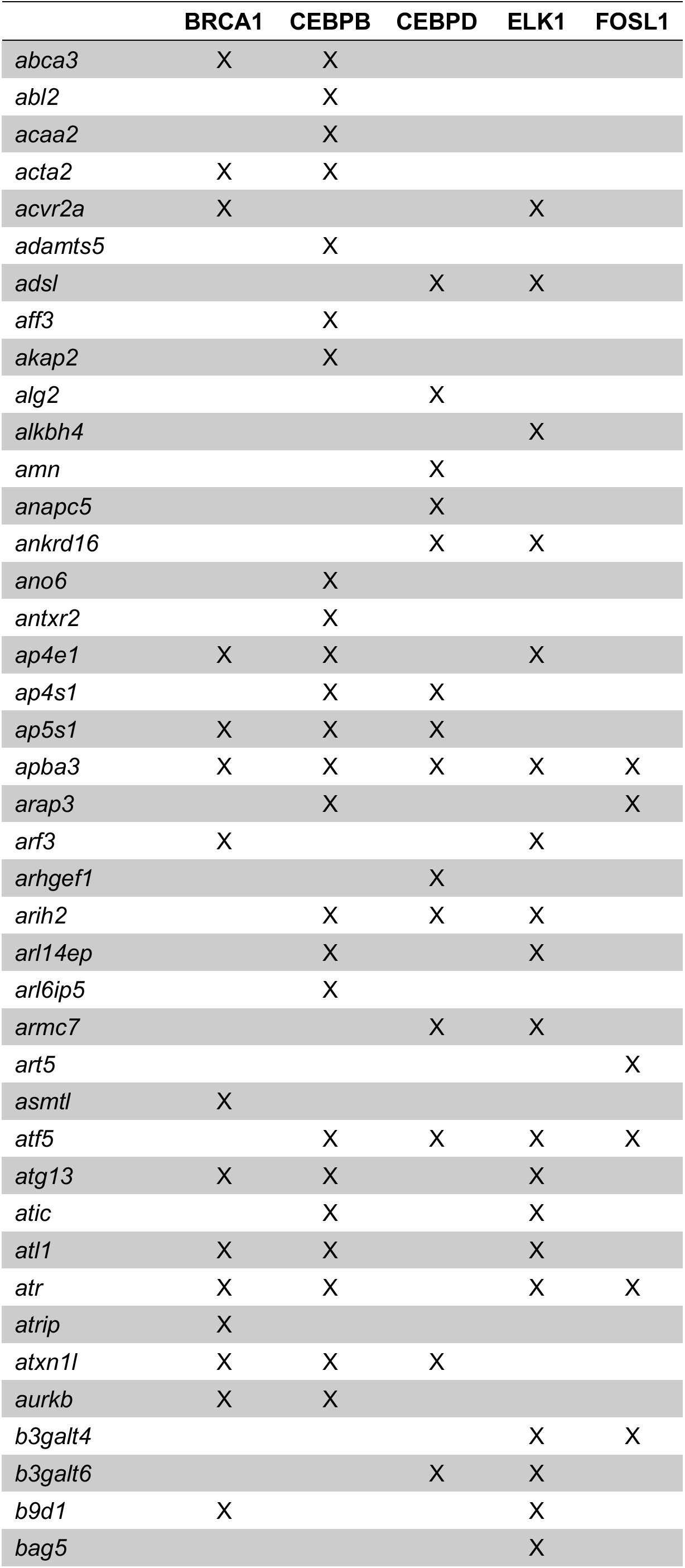
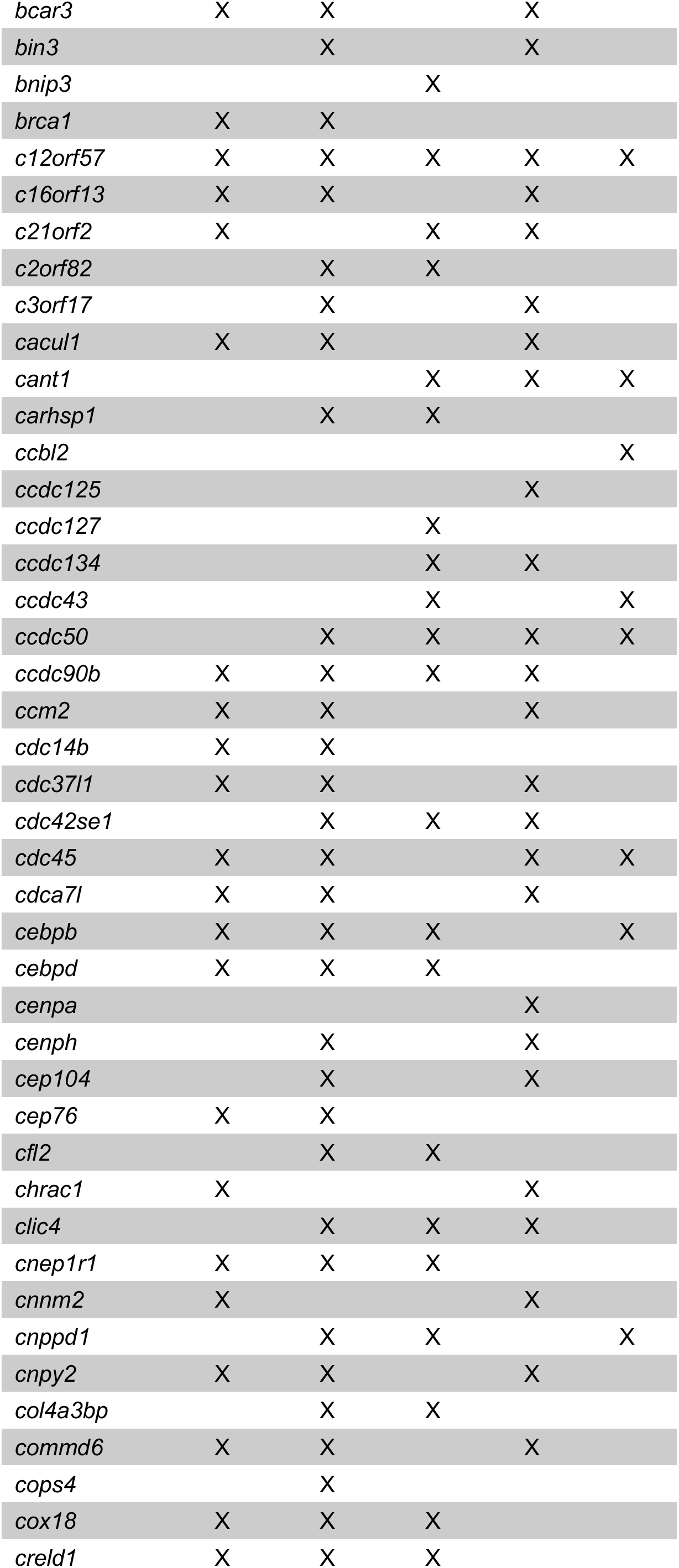
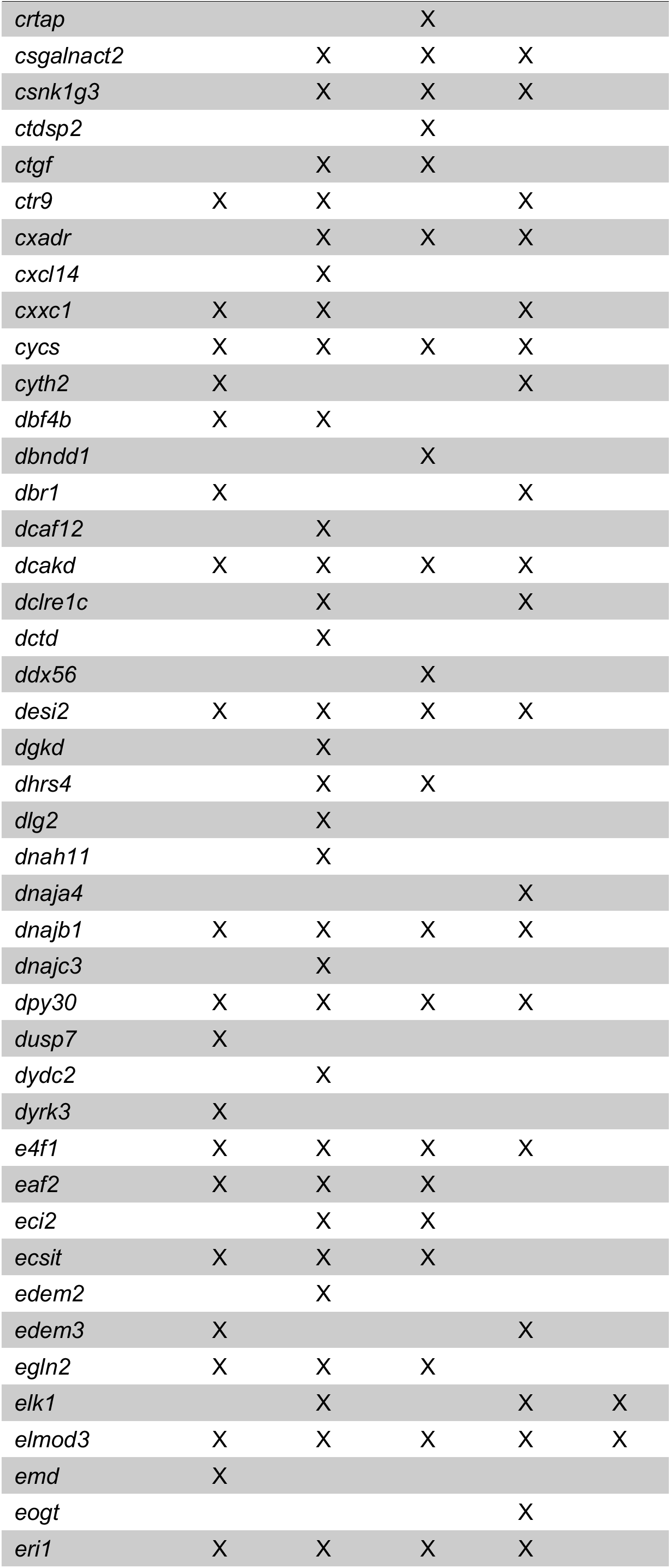
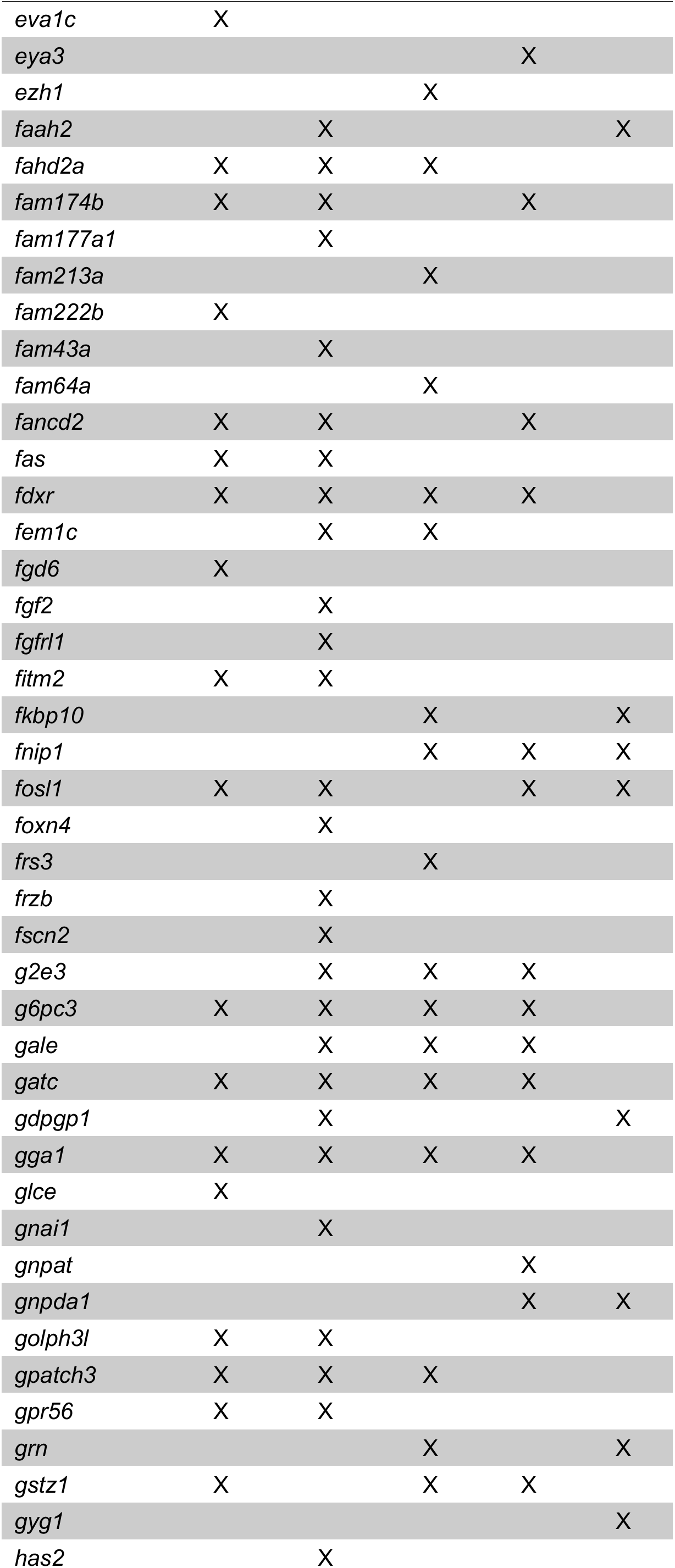
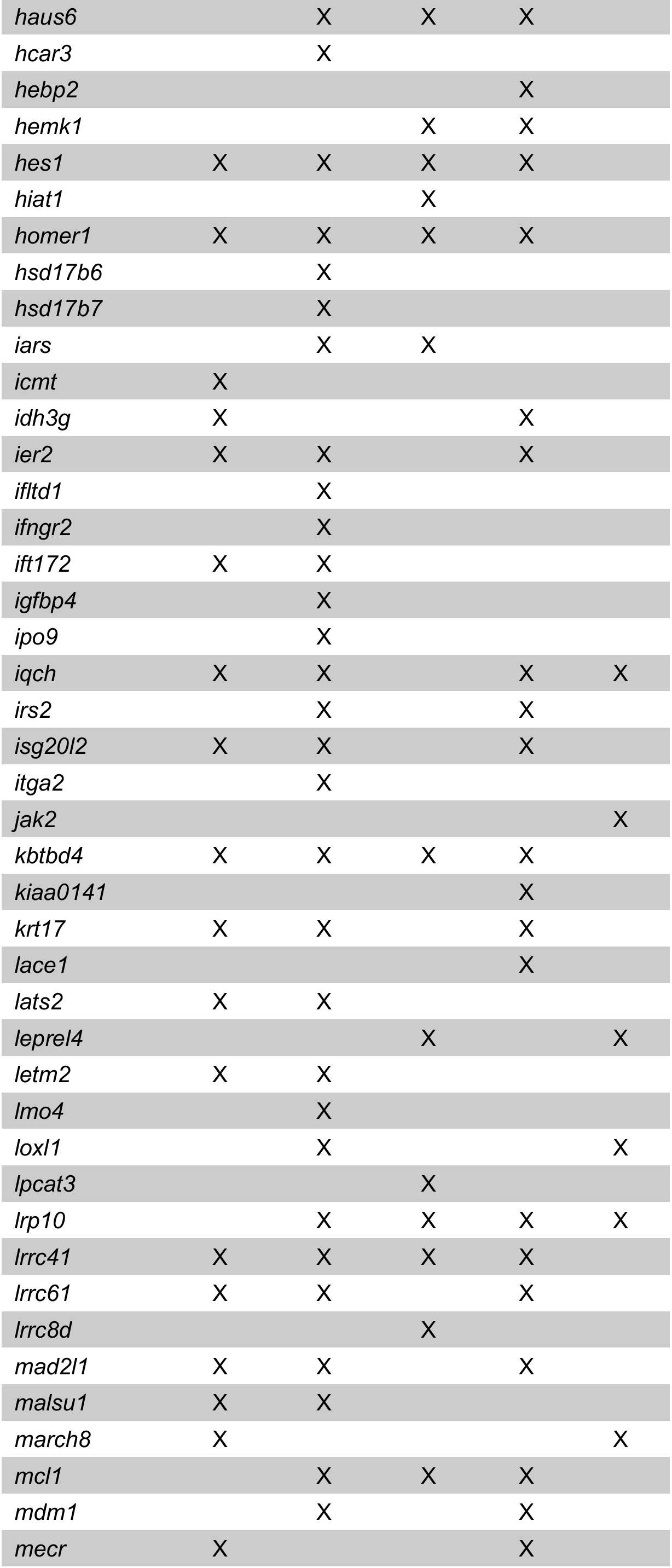
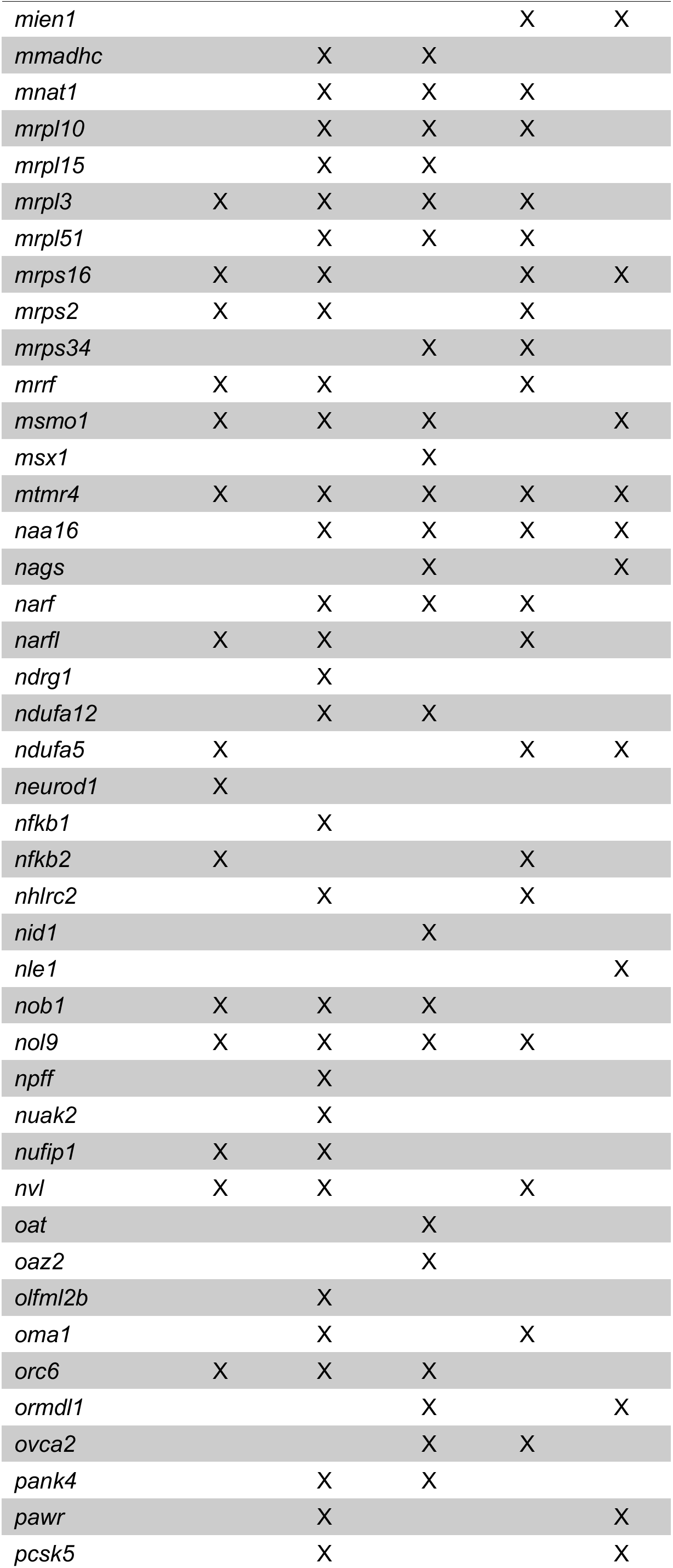
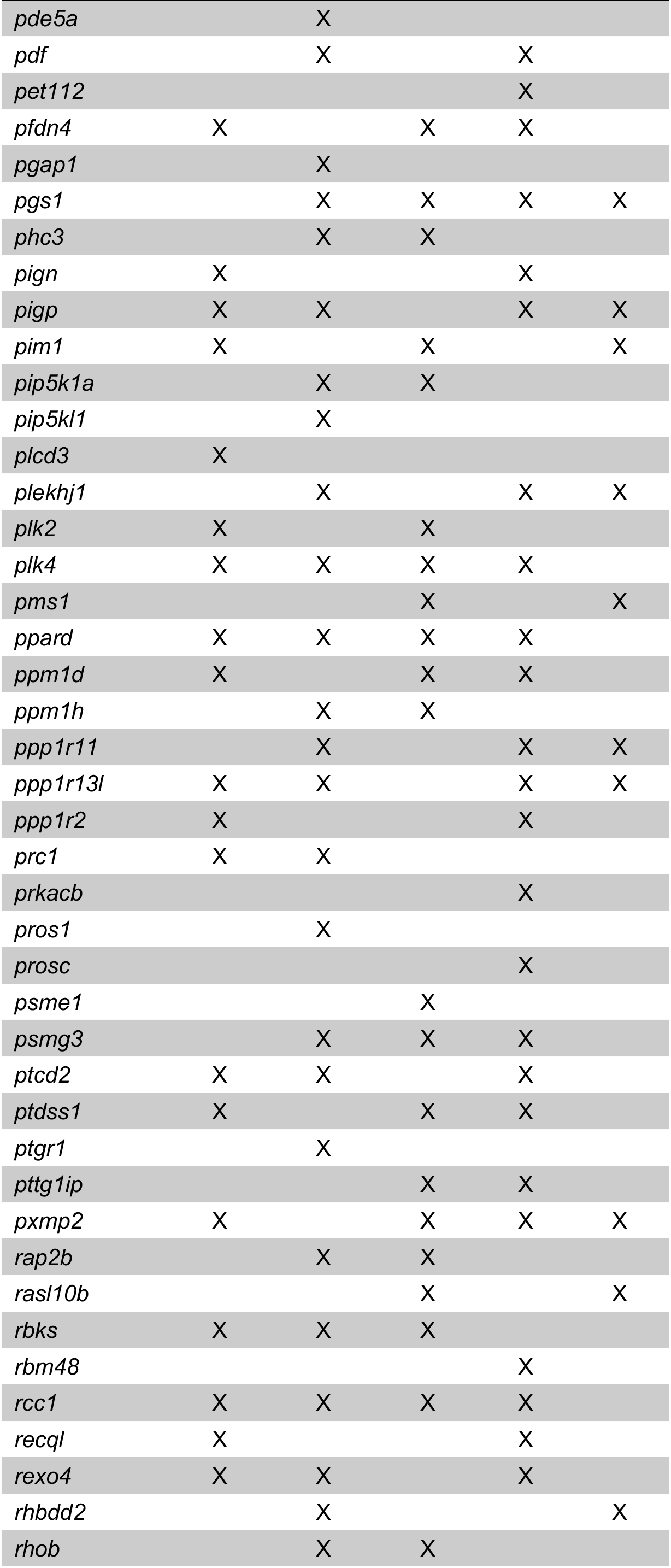
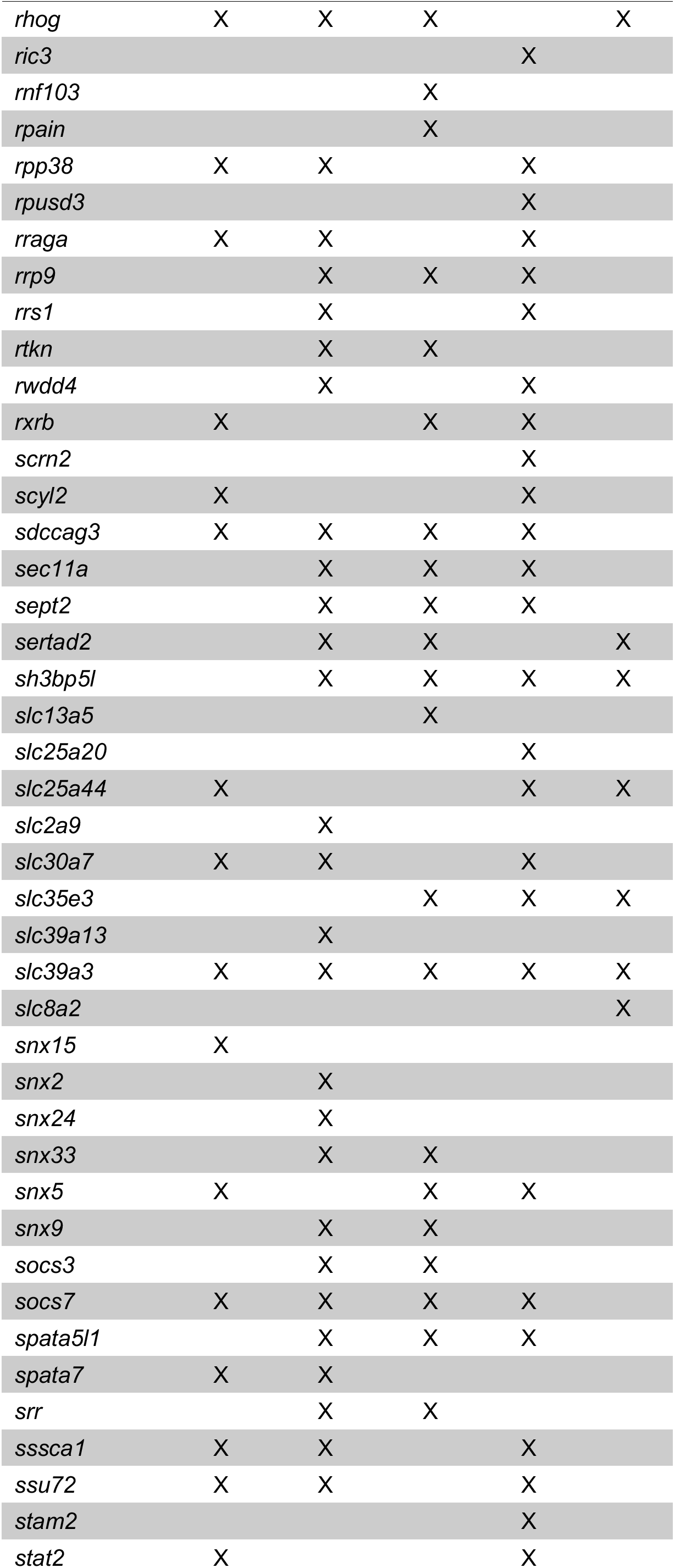
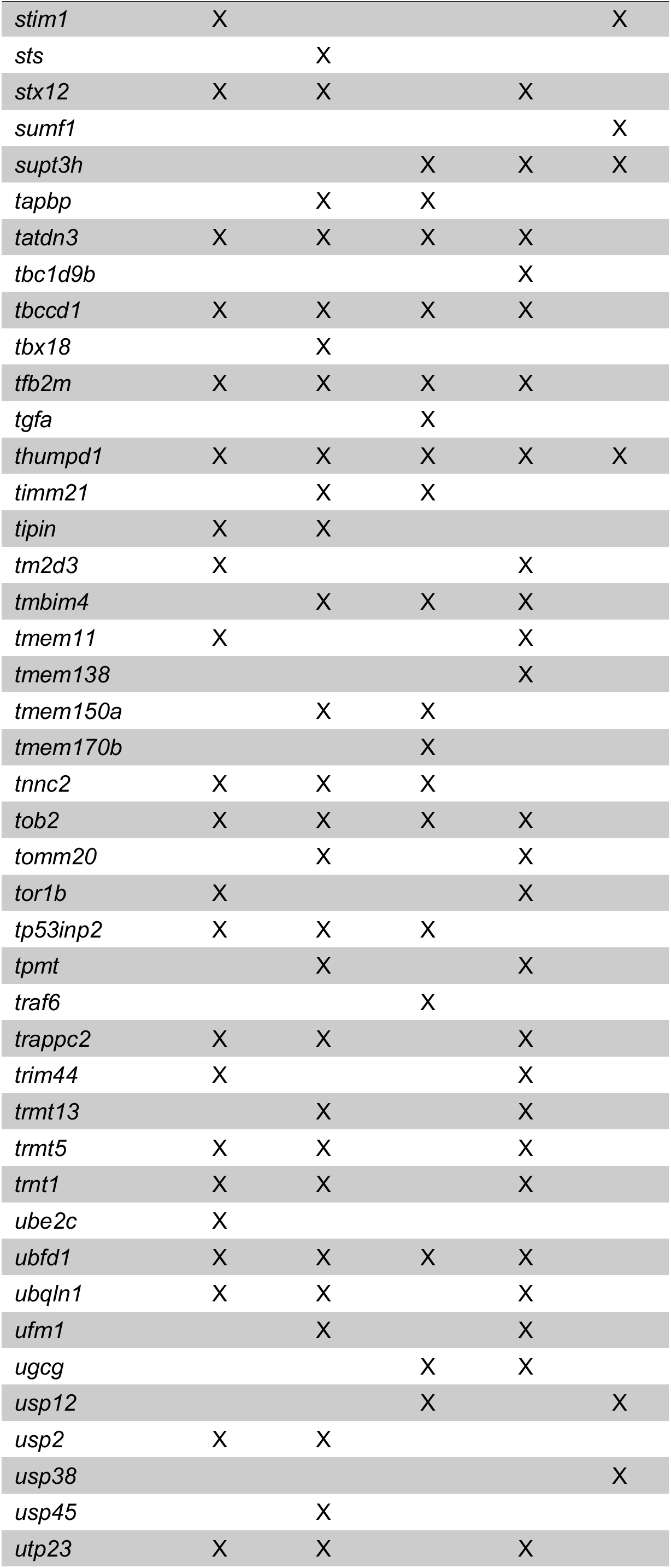
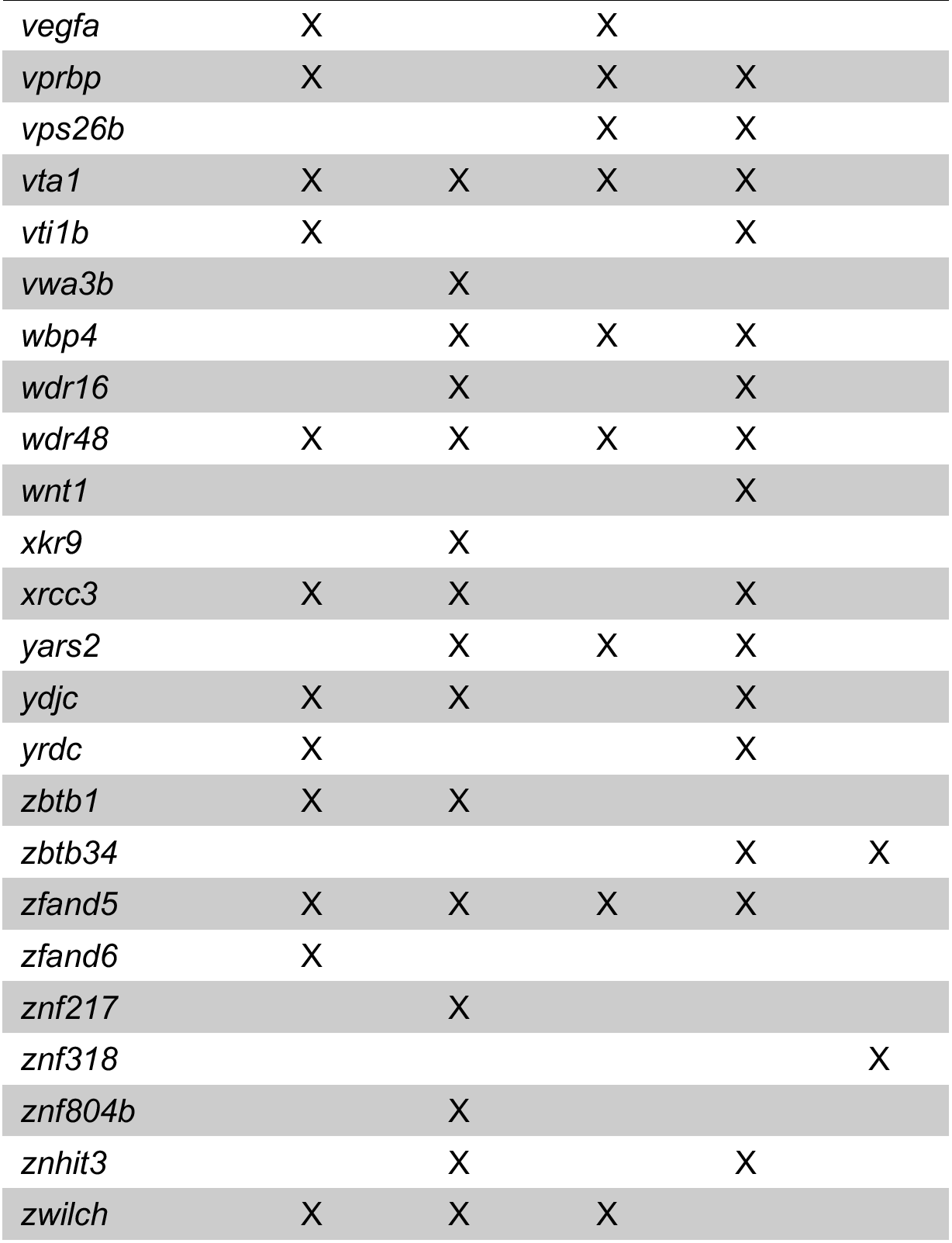
Gene list of transcripts regulated by the 5 networked transcription factors shown in Venn Diagram in Figure 4C.

**Supplemental Data S1.** Total list of DE transcripts.

**Supplemental Data S2.** Corresponding to Figure 2A (list of genes in PANTHER GO categories)

**Supplemental Data S3.** Corresponding to Figure 3A (string analysis of DE genes)

**Supplemental Data S4.** Corresponding to Figure 4A (ENCODE data of transcription factors)

## Notes

### Competing Interest Statement

The authors have declared no competing interest.

## References

Aizenman, C.D., and Cline, H.T. (2007). Enhanced visual activity in vivo forms nascent synapses in the developing retinotectal projection. J Neurophysiol 97, 2949–2957.

Anders, S., Pyl, P.T., and Huber, W. (2015). HTSeq--a Python framework to work with high-throughput sequencing data. Bioinformatics 31, 166–169.

Azim, K., Hurtado-Chong, A., Fischer, B., Kumar, N., Zweifel, S., Taylor, V., and Raineteau, O. (2015). Transcriptional Hallmarks of Heterogeneous Neural Stem Cell Niches of the Subventricular Zone. Stem Cells 33, 2232–2242.

Barbieri, F., Pattarozzi, A., Gatti, M., Porcile, C., Bajetto, A., Ferrari, A., Culler, M.D., and Florio, T. (2008). Somatostatin receptors 1, 2, and 5 cooperate in the somatostatin inhibition of C6 glioma cell proliferation in vitro via a phosphotyrosine phosphatase-eta-dependent inhibition of extracellularly regulated kinase-1/2. Endocrinology 149, 4736–4746.

Berger, C., Harzer, H., Burkard, T.R., Steinmann, J., van der Horst, S., Laurenson, A.S., Novatchkova, M., Reichert, H., and Knoblich, J.A. (2012). FACS purification and transcriptome analysis of drosophila neural stem cells reveals a role for Klumpfuss in self-renewal. Cell Rep 2, 407–418.

Besnard, A., Galan-Rodriguez, B., Vanhoutte, P., and Caboche, J. (2011). Elk-1 a transcription factor with multiple facets in the brain. Front Neurosci 5, 35.

Bestman, J.E., and Cline, H.T. (2008). The RNA binding protein CPEB regulates dendrite morphogenesis and neuronal circuit assembly in vivo. Proc Natl Acad Sci U S A 105, 20494–20499.

Bestman, J.E., Huang, L.C., Lee-Osbourne, J., Cheung, P., and Cline, H.T. (2015). An in vivo screen to identify candidate neurogenic genes in the developing Xenopus visual system. Dev Biol 408, 269–291.

Bestman, J.E., Lee-Osbourne, J., and Cline, H.T. (2012). In vivo time-lapse imaging of cell proliferation and differentiation in the optic tectum of Xenopus laevis tadpoles. J Comp Neurol 520, 401–433.

Bindea, G., Mlecnik, B., Hackl, H., Charoentong, P., Tosolini, M., Kirilovsky, A., Fridman, W.H., Pages, F., Trajanoski, Z., and Galon, J. (2009). ClueGO: a Cytoscape plug-in to decipher functionally grouped gene ontology and pathway annotation networks. Bioinformatics 25, 1091–1093.

Bolger, A.M., Lohse, M., and Usadel, B. (2014). Trimmomatic: a flexible trimmer for Illumina sequence data. Bioinformatics 30, 2114–2120.

Cavanagh, J.F., Mione, M.C., Pappas, I.S., and Parnavelas, J.G. (1997). Basic fibroblast growth factor prolongs the proliferation of rat cortical progenitor cells in vitro without altering their cell cycle parameters. Cereb Cortex 7, 293–302.

Chai, Y., Chipitsyna, G., Cui, J., Liao, B., Liu, S., Aysola, K., Yezdani, M., Reddy, E.S., and Rao, V.N. (2001). c-Fos oncogene regulator Elk-1 interacts with BRCA1 splice variants BRCA1a/1b and enhances BRCA1a/1b-mediated growth suppression in breast cancer cells. Oncogene 20, 1357–1367.

Chen, Y., Wang, H., Yoon, S.O., Xu, X., Hottiger, M.O., Svaren, J., Nave, K.A., Kim, H.A., Olson, E.N., and Lu, Q.R. (2011). HDAC-mediated deacetylation of NF-kappaB is critical for Schwann cell myelination. Nat Neurosci 14, 437–441.

Chen, Y.L., Monteith, N., Law, P.Y., and Loh, H.H. (2010). Dynamic association of p300 with the promoter of the G protein-coupled rat delta opioid receptor gene during NGF-induced neuronal differentiation. Biochem Biophys Res Commun 396, 294–298.

Chenn, A., and Walsh, C.A. (2002). Regulation of cerebral cortical size by control of cell cycle exit in neural precursors. Science 297, 365–369.

Chung, H., Li, E., Kim, Y., Kim, S., and Park, S. (2013). Multiple signaling pathways mediate ghrelin-induced proliferation of hippocampal neural stem cells. J Endocrinol 218, 49–59.

Collins, L.M. (2018). Optimization of behavioral, biobehavioral, and biomedical interventions: The multiphase optimization strategy (MOST). (New York: Springer).

Consortium, E.P. (2012). An integrated encyclopedia of DNA elements in the human genome. Nature 489, 57–74.

Cowger, J.J., Zhao, Q., Isovic, M., and Torchia, J. (2007). Biochemical characterization of the zinc-finger protein 217 transcriptional repressor complex: identification of a ZNF217 consensus recognition sequence. Oncogene 26, 3378–3386.

Denis-Donini, S., Dellarole, A., Crociara, P., Francese, M.T., Bortolotto, V., Quadrato, G., Canonico, P.L., Orsetti, M., Ghi, P., Memo, M., et al. (2008). Impaired adult neurogenesis associated with short-term memory defects in NF-kappaB p50-deficient mice. J Neurosci 28, 3911–3919.

Densham, R.M., and Morris, J.R. (2017). The BRCA1 Ubiquitin ligase function sets a new trend for remodelling in DNA repair. Nucleus 8, 116–125.

Dobin, A., Davis, C.A., Schlesinger, F., Drenkow, J., Zaleski, C., Jha, S., Batut, P., Chaisson, M., and Gingeras, T.R. (2013). STAR: ultrafast universal RNA-seq aligner. Bioinformatics 29, 15–21.

Faulkner, R.L., Wishard, T.J., Thompson, C.K., Liu, H.H., and Cline, H.T. (2015). FMRP regulates neurogenesis in vivo in Xenopus laevis tadpoles. eNeuro 2, e0055.

Frappart, P.O., Lee, Y., Lamont, J., and McKinnon, P.J. (2007). BRCA2 is required for neurogenesis and suppression of medulloblastoma. EMBO J 26, 2732–2742.

Gaete, M., Munoz, R., Sanchez, N., Tampe, R., Moreno, M., Contreras, E.G., Lee-Liu, D., and Larrain, J. (2012). Spinal cord regeneration in Xenopus tadpoles proceeds through activation of Sox2-positive cells. Neural Dev 7, 13.

Gambrill, A.C., Faulkner, R.L., McKeown, C.R., and Cline, H.T. (2019). Enhanced visual experience rehabilitates the injured brain in Xenopus tadpoles in an NMDAR-dependent manner. J Neurophysiol 121, 306–320.

Gao, B., Shen, X., Kunos, G., Meng, Q., Goldberg, I.D., Rosen, E.M., and Fan, S. (2001). Constitutive activation of JAK-STAT3 signaling by BRCA1 in human prostate cancer cells. FEBS Lett 488, 179–184.

Gearhart, J., Pashos, E.E., and Prasad, M.K. (2007). Pluripotency redux--advances in stem-cell research. N Engl J Med 357, 1469–1472.

Goertsches, R., Comabella, M., Navarro, A., Perkal, H., and Montalban, X. (2005). Genetic association between polymorphisms in the ADAMTS14 gene and multiple sclerosis. J Neuroimmunol 164, 140–147.

Gotz, M., and Huttner, W.B. (2005). The cell biology of neurogenesis. Nat Rev Mol Cell Biol 6, 777–788.

Gotz, M., Nakafuku, M., and Petrik, D. (2016). Neurogenesis in the Developing and Adult Brain-Similarities and Key Differences. Cold Spring Harb Perspect Biol 8.

Gowen, L.C., Johnson, B.L., Latour, A.M., Sulik, K.K., and Koller, B.H. (1996). Brca1 deficiency results in early embryonic lethality characterized by neuroepithelial abnormalities. Nat Genet 12, 191–194.

Hall, Z.J., and Tropepe, V. (2018). Movement maintains forebrain neurogenesis via peripheral neural feedback in larval zebrafish. Elife 7.

Hamel, M.G., Ajmo, J.M., Leonardo, C.C., Zuo, F., Sandy, J.D., and Gottschall, P.E. (2008). Multimodal signaling by the ADAMTSs (a disintegrin and metalloproteinase with thrombospondin motifs) promotes neurite extension. Exp Neurol 210, 428–440.

Hanlon, M., Bundy, L.M., and Sealy, L. (2000). C/EBP beta and Elk-1 synergistically transactivate the c-fos serum response element. BMC Cell Biol 1, 2.

He, J.Y., Xi, W.H., Zhu, L.B., Long, X.H., Chen, X.Y., Liu, J.M., Luo, Q.F., Zhu, X.P., and Liu, Z.L. (2015). Knockdown of Aurora-B alters osteosarcoma cell malignant phenotype via decreasing phosphorylation of VCP and NF-kappaB signaling. Tumour Biol 36, 3895–3902.

Hess, J., Angel, P., and Schorpp-Kistner, M. (2004). AP-1 subunits: quarrel and harmony among siblings. J Cell Sci 117, 5965–5973.

Hu, W.F., Chahrour, M.H., and Walsh, C.A. (2014). The diverse genetic landscape of neurodevelopmental disorders. Annual review of genomics and human genetics 15, 195–213.

Jia, Z., Gao, S., M’Rabet, N., De Geyter, C., and Zhang, H. (2014). Sp1 is necessary for gene activation of Adamts17 by estrogen. J Cell Biochem 115, 1829–1839.

Joukov, V., Chen, J., Fox, E.A., Green, J.B., and Livingston, D.M. (2001). Functional communication between endogenous BRCA1 and its partner, BARD1, during Xenopus laevis development. Proc Natl Acad Sci U S A 98, 12078–12083.

Kakebeen, A.D., Chitsazan, A.D., Williams, M.C., Saunders, L.M., and Wills, A.E. (2020). Chromatin accessibility dynamics and single cell RNA-Seq reveal new regulators of regeneration in neural progenitors. Elife 9.

Kang, M.K., and Kang, S.K. (2008). Interleukin-6 induces proliferation in adult spinal cord-derived neural progenitors via the JAK2/STAT3 pathway with EGF-induced MAPK phosphorylation. Cell Prolif 41, 377–392.

Kim, Y.H., Chung, J.I., Woo, H.G., Jung, Y.S., Lee, S.H., Moon, C.H., Suh-Kim, H., and Baik, E.J. (2010). Differential regulation of proliferation and differentiation in neural precursor cells by the Jak pathway. Stem Cells 28, 1816–1828.

Kirischuk, S., Sinning, A., Blanquie, O., Yang, J.W., Luhmann, H.J., and Kilb, W. (2017). Modulation of Neocortical Development by Early Neuronal Activity: Physiology and Pathophysiology. Front Cell Neurosci 11, 379.

Lassmann, T., Hayashizaki, Y., and Daub, C.O. (2011). SAMStat: monitoring biases in next generation sequencing data. Bioinformatics 27, 130–131.

Li, H.Y., Liao, C.Y., Lee, K.H., Chang, H.C., Chen, Y.J., Chao, K.C., Chang, S.P., Cheng, H.Y., Chang, C.M., Chang, Y.L., et al. (2011). Collagen IV significantly enhances migration and transplantation of embryonic stem cells: involvement of alpha2beta1 integrin-mediated actin remodeling. Cell Transplant 20, 893–907.

Louvi, A., and Artavanis-Tsakonas, S. (2006). Notch signalling in vertebrate neural development. Nat Rev Neurosci 7, 93–102.

Love, M.I., Huber, W., and Anders, S. (2014). Moderated estimation of fold change and dispersion for RNA-seq data with DESeq2. Genome Biol 15, 550.

Luhmann, H.J., Sinning, A., Yang, J.W., Reyes-Puerta, V., Stuttgen, M.C., Kirischuk, S., and Kilb, W. (2016). Spontaneous Neuronal Activity in Developing Neocortical Networks: From Single Cells to Large-Scale Interactions. Front Neural Circuits 10, 40.

Luo, J., Su, F., Chen, D., Shiloh, A., and Gu, W. (2000). Deacetylation of p53 modulates its effect on cell growth and apoptosis. Nature 408, 377–381.

Ma, D.K., Jang, M.-H., Guo, J.U., Kitabatake, Y., Chang, M.-l., Pow-anpongkul, N., Flavell, R.A., Lu, B., G-l., M., and Song, H. (2009). Neuronal Activity–Induced Gadd45b Promotes Epigenetic DNA Demethylation and Adult Neurogenesis. Science 323, 1074–1077.

Madsen, T.M., Treschow, A., Bengzon, J., Bolwig, T.G., Lindvall, O., and Tingstrom, A. (2000). Increased neurogenesis in a model of electroconvulsive therapy. Biol Psychiatry 47, 1043–1049.

Maniccia, A.W., Lewis, C., Begum, N., Xu, J., Cui, J., Chipitsyna, G., Aysola, K., Reddy, V., Bhat, G., Fujimura, Y., et al. (2009). Mitochondrial localization, ELK-1 transcriptional regulation and growth inhibitory functions of BRCA1, BRCA1a, and BRCA1b proteins. J Cell Physiol 219, 634–641.

Marei, H.E., Althani, A., Afifi, N., Michetti, F., Pescatori, M., Pallini, R., Casalbore, P., Cenciarelli, C., Schwartz, P., and Ahmed, A.E. (2011). Gene expression profiling of embryonic human neural stem cells and dopaminergic neurons from adult human substantia nigra. PLoS One 6, e28420.

Mi, H., Muruganujan, A., and Thomas, P.D. (2013). PANTHER in 2013: modeling the evolution of gene function, and other gene attributes, in the context of phylogenetic trees. Nucleic Acids Res 41, D377–386.

Mullan, P.B., Quinn, J.E., and Harkin, D.P. (2006). The role of BRCA1 in transcriptional regulation and cell cycle control. Oncogene 25, 5854–5863.

Nagel, M., Tahinci, E., Symes, K., and Winklbauer, R. (2004). Guidance of mesoderm cell migration in the Xenopus gastrula requires PDGF signaling. Development 131, 2727–2736.

Orii, K.E., Lee, Y., Kondo, N., and McKinnon, P.J. (2006). Selective utilization of nonhomologous end-joining and homologous recombination DNA repair pathways during nervous system development. Proc Natl Acad Sci U S A 103, 10017–10022.

Pan, Y., and Monje, M. (2020). Activity Shapes Neural Circuit Form and Function: A Historical Perspective. J Neurosci 40, 944–954.

Pao, G.M., Zhu, Q., Perez-Garcia, C.G., Chou, S.J., Suh, H., Gage, F.H., O’Leary, D.D., and Verma, I.M. (2014). Role of BRCA1 in brain development. Proc Natl Acad Sci U S A 111, E1240–1248.

Pataskar, A., Jung, J., Smialowski, P., Noack, F., Calegari, F., Straub, T., and Tiwari, V.K. (2015). NeuroD1 reprograms chromatin and transcription factor landscapes to induce the neuronal program. EMBO J.

Paul, C., Lacroix, M., Iankova, I., Julien, E., Schafer, B.W., Labalette, C., Wei, Y., Le Cam, A., Le Cam, L., and Sardet, C. (2006). The LIM-only protein FHL2 is a negative regulator of E4F1. Oncogene 25, 5475–5484.

Pennanen, P.T., Sarvilinna, N.S., Toimela, T., and Ylikomi, T.J. (2011). Inhibition of FOSL1 overexpression in antiestrogen-resistant MCF-7 cells decreases cell growth and increases vacuolization and cell death. Steroids 76, 1063–1068.

Pires-daSilva, A., and Sommer, R.J. (2003). The evolution of signalling pathways in animal development. Nat Rev Genet 4, 39–49.

Prakash, R., Zhang, Y., Feng, W., and Jasin, M. (2015). Homologous recombination and human health: the roles of BRCA1, BRCA2, and associated proteins. Cold Spring Harb Perspect Biol 7, a016600.

Pramparo, T., Lombardo, M.V., Campbell, K., Barnes, C.C., Marinero, S., Solso, S., Young, J., Mayo, M., Dale, A., Ahrens-Barbeau, C., et al. (2015). Cell cycle networks link gene expression dysregulation, mutation, and brain maldevelopment in autistic toddlers. Mol Syst Biol 11, 841.

Pulido-Salgado, M., Vidal-Taboada, J.M., and Saura, J. (2015). C/EBPbeta and C/EBPdelta transcription factors: Basic biology and roles in the CNS. Prog Neurobiol 132, 1–33.

Pulvers, J.N., and Huttner, W.B. (2009). Brca1 is required for embryonic development of the mouse cerebral cortex to normal size by preventing apoptosis of early neural progenitors. Development 136, 1859–1868.

Qiao, Y., Wang, R., Yang, X., Tang, K., and Jing, N. (2015). Dual roles of histone H3 lysine 9 acetylation in human embryonic stem cell pluripotency and neural differentiation. J Biol Chem 290, 2508–2520.

Romero-Calvo, I., Ocon, B., Martinez-Moya, P., Suarez, M.D., Zarzuelo, A., Martinez-Augustin, O., and de Medina, F.S. (2010). Reversible Ponceau staining as a loading control alternative to actin in Western blots. Anal Biochem 401, 318–320.

Rosenstein, J.M., Mani, N., Khaibullina, A., and Krum, J.M. (2003). Neurotrophic effects of vascular endothelial growth factor on organotypic cortical explants and primary cortical neurons. J Neurosci 23, 11036–11044.

Ruthazer, E.S., Li, J., and Cline, H.T. (2006). Stabilization of axon branch dynamics by synaptic maturation. J Neurosci 26, 3594–3603.

Sanalkumar, R., Vidyanand, S., Lalitha Indulekha, C., and James, J. (2010). Neuronal vs. glial fate of embryonic stem cell-derived neural progenitors (ES-NPs) is determined by FGF2/EGF during proliferation. J Mol Neurosci 42, 17–27.

Schindelin, J., Arganda-Carreras, I., Frise, E., Kaynig, V., Longair, M., Pietzsch, T., Preibisch, S., Rueden, C., Saalfeld, S., Schmid, B., et al. (2012). Fiji: an open-source platform for biological-image analysis. Nat Methods 9, 676–682.

Schneider, C.A., Rasband, W.S., and Eliceiri, K.W. (2012). NIH Image to ImageJ: 25 years of image analysis. Nat Methods 9, 671–675.

Schreiber, M., Kolbus, A., Piu, F., Szabowski, A., Mohle-Steinlein, U., Tian, J., Karin, M., Angel, P., and Wagner, E.F. (1999). Control of cell cycle progression by c-Jun is p53 dependent. Genes Dev 13, 607–619.

Sharma, P., and Cline, H.T. (2010). Visual activity regulates neural progenitor cells in developing xenopus CNS through musashi1. Neuron 68, 442–455.

Sierra, A., Martin-Suarez, S., Valcarcel-Martin, R., Pascual-Brazo, J., Aelvoet, S.A., Abiega, O., Deudero, J.J., Brewster, A.L., Bernales, I., Anderson, A.E., et al. (2015). Neuronal hyperactivity accelerates depletion of neural stem cells and impairs hippocampal neurogenesis. Cell Stem Cell 16, 488–503.

Sin, W.C., Haas, K., Ruthazer, E.S., and Cline, H.T. (2002). Dendrite growth increased by visual activity requires NMDA receptor and Rho GTPases. Nature 419, 475–480.

Sogut, M.S., Venugopal, C., Kandemir, B., Dag, U., Mahendram, S., Singh, S., Gulfidan, G., Arga, K.Y., Yilmaz, B., and Kurnaz, I.A. (2021). ETS-Domain Transcription Factor Elk-1 Regulates Stemness Genes in Brain Tumors and CD133+ BrainTumor-Initiating Cells. J Pers Med 11.

Song, J., Zhong, C., Bonaguidi, M.A., Sun, G.J., Hsu, D., Gu, Y., Meletis, K., Huang, Z.J., Ge, S., Enikolopov, G., et al. (2012). Neuronal circuitry mechanism regulating adult quiescent neural stem-cell fate decision. Nature 489, 150–154.

Szklarczyk, D., Franceschini, A., Wyder, S., Forslund, K., Heller, D., Huerta-Cepas, J., Simonovic, M., Roth, A., Santos, A., Tsafou, K.P., et al. (2015). STRING v10: protein-protein interaction networks, integrated over the tree of life. Nucleic Acids Res 43, D447–452.

Thaeron, C., Avaron, F., Casane, D., Borday, V., Thisse, B., Thisse, C., Boulekbache, H., and Laurenti, P. (2000). Zebrafish evx1 is dynamically expressed during embryogenesis in subsets of interneurones, posterior gut and urogenital system. Mech Dev 99, 167–172.

Thompson, C.K., and Cline, H.T. (2016). Thyroid Hormone Acts Locally to Increase Neurogenesis, Neuronal Differentiation, and Dendritic Arbor Elaboration in the Tadpole Visual System. J Neurosci 36, 10356–10375.

Trapnell, C., Roberts, A., Goff, L., Pertea, G., Kim, D., Kelley, D.R., Pimentel, H., Salzberg, S.L., Rinn, J.L., and Pachter, L. (2012). Differential gene and transcript expression analysis of RNA-seq experiments with TopHat and Cufflinks. Nat Protoc 7, 562–578.

Tropepe, V., Craig, C.G., Morshead, C.M., and van der Kooy, D. (1997). Transforming growth factor-alpha null and senescent mice show decreased neural progenitor cell proliferation in the forebrain subependyma. J Neurosci 17, 7850–7859.

Vallejo, A., Perurena, N., Guruceaga, E., Mazur, P.K., Martinez-Canarias, S., Zandueta, C., Valencia, K., Arricibita, A., Gwinn, D., Sayles, L.C., et al. (2017). An integrative approach unveils FOSL1 as an oncogene vulnerability in KRAS-driven lung and pancreatic cancer. Nat Commun 8, 14294.

van de Leemput, J., Boles, N.C., Kiehl, T.R., Corneo, B., Lederman, P., Menon, V., Lee, C., Martinez, R.A., Levi, B.P., Thompson, C.L., et al. (2014). CORTECON: a temporal transcriptome analysis of in vitro human cerebral cortex development from human embryonic stem cells. Neuron 83, 51–68.

Wang, H., Wang, X., Qu, J., Yue, Q., Hu, Y., and Zhang, H. (2015). VEGF Enhances the Migration of MSCs in Neural Differentiation by Regulating Focal Adhesion Turnover. J Cell Physiol 230, 2728–2742.

Wang, J., Zhang, H., Young, A.G., Qiu, R., Argalian, S., Li, X., Wu, X., Lemke, G., and Lu, Q. (2011). Transcriptome analysis of neural progenitor cells by a genetic dual reporter strategy. Stem Cells 29, 1589–1600.

Wang, S., and Zhao, J. (2015). Multi-attribute intergrated measurement of node importance in complex networks. Chaos 25.

Wells, T., Rough, K., and Carter, D.A. (2011). Transcription Mapping of Embryonic Rat Brain Reveals EGR-1 Induction in SOX2 Neural Progenitor Cells. Front Mol Neurosci 4, 6.

Yang, Y., Shen, W., Ni, Y., Su, Y., Yang, Z., and Zhao, C. (2015). Impaired Interneuron Development after Foxg1 Disruption. Cereb Cortex.

Yap, E.L., and Greenberg, M.E. (2018). Activity-Regulated Transcription: Bridging the Gap between Neural Activity and Behavior. Neuron 100, 330–348.

Zhang, J., Fei, T., Li, Z., Zhu, G., Wang, L., and Chen, Y.G. (2013). BMP induces cochlin expression to facilitate self-renewal and suppress neural differentiation of mouse embryonic stem cells. J Biol Chem 288, 8053–8060.

Zheng, H., Ying, H., Yan, H., Kimmelman, A.C., Hiller, D.J., Chen, A.J., Perry, S.R., Tonon, G., Chu, G.C., Ding, Z., et al. (2008). p53 and Pten control neural and glioma stem/progenitor cell renewal and differentiation. Nature 455, 1129–1133.

Zupanc, G.K.H. (2021). Adult neurogenesis in the central nervous system of teleost fish: from stem cells to function and evolution. J Exp Biol 224.

